# Synchronized LFP rhythmicity in the social brain reflects the context of social encounters

**DOI:** 10.1101/2023.04.26.538368

**Authors:** Alok Nath Mohapatra, David Peles, Shai Netser, Shlomo Wagner

**Affiliations:** Sagol Department of Neurobiology, Faculty of Natural Sciences, University of Haifa, Haifa, Israel

**Author notes:** Corresponding author: Alok Nath Mohapatra, Sagol Department of Neurobiology, Faculty of Natural Sciences, University of Haifa, Mt. Carmel, Haifa 3498838, Israel.

**Keywords:** social context, social discrimination, social brain, theta rhythmicity, in vivo electrophysiology, local field potential

## Abstract

Mammalian social behavior is highly context-sensitive. Yet, little is known about the mechanisms that modulate social behavior according to its context. Recent studies have revealed a network of mostly limbic brain regions, here termed the “social brain”, which regulates social behavior. We hypothesized that coherent theta and gamma rhythms reflect the organization of the social brain regions into functional networks in a context-dependent manner. To test this concept, we simultaneously recorded extracellular activity from multiple social brain regions in mice performing three social discrimination tasks. Local field potential (LFP) rhythmicity across all tasks was dominated by a general internal state. However, during stimulus investigation LFP rhythmicity was sensitive to stimulus characteristics. Specifically, the pattern of LFP coherence between the various regions reflected mainly the social context. Moreover, we found the ventral dentate gyrus to play a pivotal role in coordinating the context-specific rhythmic activity in the network.

## Introduction

Mammalian social behavior is highly complex and dynamic, involving multiple types of distinct, sometimes even opposing, interactions between partners. Indeed, the identity of a partner can completely change the nature and trajectory of social actions taken by an individual [1]. In addition to these complexities, social interactions are highly dependent upon the social context. For example, humans will most likely respond differently to a hand placed upon their shoulder from behind if this happens in a frightening context, such as in a dark alley in a foreign city, then if the same contact occurs in a cocktail party. Presently, little is known of the brain mechanisms and neural circuits that encode the context of social encounters and change responses to social cues accordingly.

In the last two decades, studies have begun to reveal the brain circuits that sub-serve various types of social behavior [for recent review papers see for example 2, 3-5]. Such studies exposed the involvement of a vast network of limbic brain regions, here termed the “social brain” [6], in processing social sensory cues and regulating mammalian social behavior [7, 8]. These include striatal regions, such as the nucleus accumbens core (AcbC) and shell (AcbSh), the prelimbic (PrL) and infralimbic (IL) prefrontal cortical areas, several hippocampal and septal areas and multiple amygdaloid and hypothalamic nuclei [9–12]. Many of these areas are highly interconnected in a bidirectional manner [13–17], and some were shown to be involved in various, at times opposing, types of social behavior [9, 18–21]. It remains, however, unclear how this intricate network of brain areas generates the large repertoire of distinct types of social behavior. Recent studies using multi-site brain recordings from behaving animals have demonstrated that system-level neural activity in sub-networks of the social brain predicts individual social preferences [22] and decision-making [23] better than does local neural activity at any single brain region. These results thus suggest that coding of the various aspects of social behavior in the brain should be considered at the system level.

Oscillatory neural activity, mostly in the theta (4-12 Hz) and gamma (30-80 Hz) bands, was reported in many cortical and sub-cortical brain regions in various species [24–26], with its power being shown to intensify during demanding cognitive functions, such as learning [27–29] and social communication [30–32]. Furthermore, abnormal theta and gamma rhythms have been reported in multiple neurodevelopmental disorders [33–35], such as autism spectrum disorder (ASD) [36, 37]. Accordingly, one prominent hypothesis states that coherent manifestation of these rhythms can dynamically coordinate the activity of neural ensembles dispersed over multiple brain regions and link them into *ad hoc* functional networks [38].

In the present study, we hypothesized that coherent theta and gamma rhythms couple various regions of the social brain into functional networks in a social context-dependent manner. In other words, distinct social contexts dictate different patterns of coordinated rhythmic activity of dispersed social brain neuronal ensembles, which in turn sub-serve context-dependent processing of social cues and consequent behavioral responses. To test this hypothesis, we recorded extracellular electrical activity simultaneously from multiple regions of the social brain in mice performing three distinct binary social discrimination tasks (contexts). The same type of social stimulus served as either preferred or less-preferred stimulus in all three tasks. Using this design, we could link distinct patterns of rhythmic neural activity across the social brain to either stimulus identity or its valence, or to the social context. Our results reveal that the pattern of coordinated oscillatory activity (coherence) in the network is strongly correlated with the social context and carries information that may be used to discriminate between distinct, albeit similar, social contexts. Further, we revealed that the ventral dentate gyrus (vDG), an area previously linked to contextual information [39, 40], seems to be involved in coordinating the coherent activity among the various regions of the social brain.

## Results

### Analyzing the behavior of CD1 male mice during three distinct binary social discrimination tasks

Using custom-built electrode arrays (EAr) we simultaneously acquired local field potential (LFP) signals from up to 16 brain regions at a time (cumulative count: 18 regions; Figure S1A and Table S1) during interactions of adult male mice (subjects; n=14) with various stimuli [41]. We aimed to sample widespread social-behavior associated regions in the cortex (prefrontal and piriform), striatum (nucleus accumbens and ventral pallidum), hippocampus (e.g. dentate gyrus and CA1), septal nuclei (latera septum), amygdala (e.g. basolateral and medial) and hypothalamus (e.g. dorsomedial and paraventricular nuclei). The location of each electrode was verified *post mortem* [41], and since the targeting accuracy was limited, not all brain regions were recorded in each subject (see Fig. S1A and Table S1 for details). For social contexts,, we employed three distinct binary social discrimination tasks [46, 47], namely the social preference (SP), (Fig 1A), emotional-state preference (EsP) (Fig. 1E) and sex preference (SxP) (Fig 1I) tasks [46]. (See timeline in Fig. S1B). Each task comprised a five min-long baseline period involving empty chambers located at opposite corners of the arena, followed by a five min-long encounter period, when a distinct stimulus was introduced into each chamber [42]. Mice performing the SP task tended to interact with social stimuli (conspecifics) for significantly more time than with objects throughout the encounter period (Fig. 1B-D). Similarly, mice performing the EsP task preferred to interact with socially isolated rather than group-housed stimuli (Fig. 1E-H), while mice performing the SxP task tended to interact more with female than with male stimuli (Fig 1I-L). Thus, in each task, the subjects discriminated between a preferred and a less-preferred stimulus. Importantly, the same type of stimulus (a group-housed male mouse) that was the preferred stimulus in the SP task was the less-preferred stimulus in the other two tasks. Therefore, this set of tasks allowed us to analyze brain-wide neural activity patterns in association with either the type of stimulus (i.e., a group-housed male vs. an object/group-housed female/isolated male) or its valence (i.e., preferred vs. less-preferred), or the social context (i.e., SP, EsP or SxP task). It should be noted, that in our hands ICR female mice do not discriminate between group-housed and isolated stimuli, hence we conducted this study using male subjects only.

**Figure 1.**
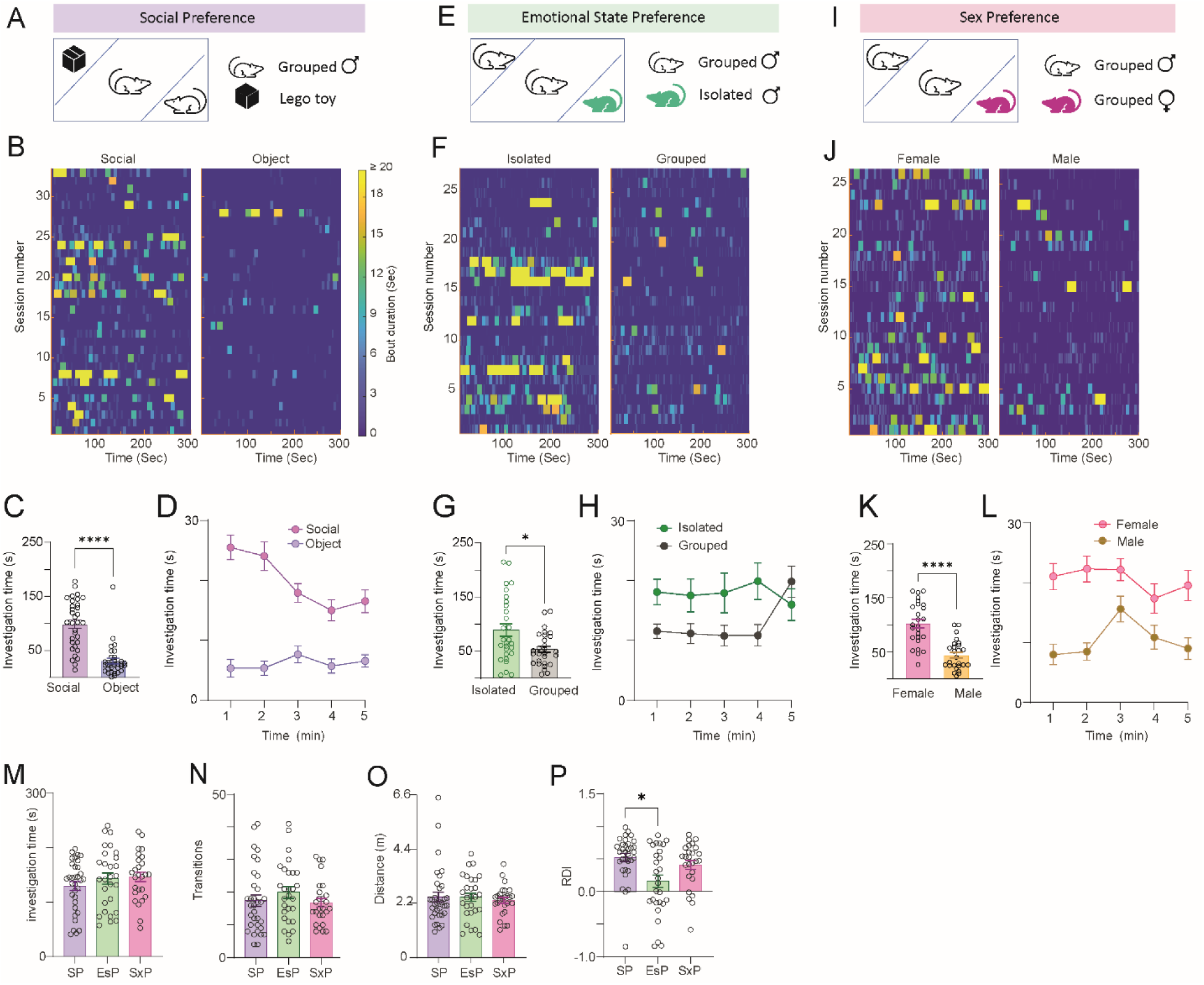
Similar behavior of subject mice across three binary social discrimination tasks. A. A scheme of the arena during SP task session. The two stimuli types are indicated on the right side. B. Heat maps of investigation bouts made by the subjects toward each of the stimuli (stimulus type is noted above) across the five min-long encounter period of the SP task, with color-coding of the investigation bout duration (see scale on the right of the panel). Each line represents a distinct session. C. Mean (±SEM) time dedicated by a subject for investigating each stimulus during the SP task sessions shown in B. Wilcoxon matched pairs signed rank test, n = 33 sessions, W = -495, \*\*\*\**p*<0.0001. D. As in C, plotted vs. time using one min bins. E-H. As in A-D, for the EsP task. Paired t-test, n = 28 sessions, t (27) =2.374, \**p* = 0.025. I-L. As in A-D, for the SxP task. Paired t-test, n = 26 sessions, t (25) =5.75, \*\*\*\**p*<0.0001. M. Mean (±SEM) total time dedicated by a subject to investigate both stimuli during the encounter stage of each task. N. Mean (±SEM) number of transitions between stimuli made by the subject during the encounter period of each task. O. Mean (±SEM) distance traveled by the subjects during the encounter stage of each task. P. Mean (±SEM) RDI for each task. Kruskal-Wallis test, n = 3 tests, 87 sessions, H = 8.509, *p* = 0.0142; Dunn’s *post-hoc* test, \**p*<0.05.

We further compared multiple behavioral parameters across the various tasks. There were no significant differences between the tasks in terms of total time dedicated to stimuli investigation (Fig. 1M), the total number of transitions made by the subjects between the two stimuli (Fig. 1N) or the distance traveled by the subjects during a task (Fig. 1O). Nonetheless, the preference (reflected by the relative discrimination index, RDI) between the two stimuli was lower in the EsP task, as compared to the SP task (Fig. 1P). Overall, subject behavior was similar across the various tasks.

### Different tasks elicit different profiles of rhythmic LFP signals in multiple brain regions

The power spectral density (PSD) profiles of LFP signals recorded during the encounter period (Fig. 2A-B), differed among the various tasks performed by the same subject in a brain region-specific manner (Fig. S1C). For quantitative comparison, we calculated the mean theta (θP) and gamma (γP) power separately for the baseline and encounter periods of each task for each brain region. While the mean power during baseline across all regions did not significantly differ between the tasks (Fig. 2C-D), the change in power during the encounter for both theta (ΔθP) and gamma (ΔγP) rhythms was highest for the SP task, compared to the other two (Fig 2E-F). Thus, despite the generally similar behavior exhibited by subjects across the tasks (Fig. 1), their system-level brain LFP signals significantly and consistently differed in power across tasks. Specifically, despite involving only one social stimulus, the SP task induced the strongest LFP rhythmicity. When considering each brain region separately, we found that in almost all cases, the mean power of both rhythms was enhanced during the encounter period, as compared to baseline. Notably, the mean power change differed significantly among the various tasks and specific regions (Fig. 2G-H). The similar patterns of ΔθP and ΔγP across the various regions suggested the existence of a link between them. Accordingly, we found statistically significant correlation (Pearson’s, r>0.25, *p*<0.05) between ΔθP and ΔγP for the SP and SxP tasks, while borderline significant correlation (r=0.45, *p*=0.059) was observed for the EsP task (Fig. 2K-M). Thus, when measured over the course of the entire session, both rhythms seem to be driven by the same process.

**Figure 2.**
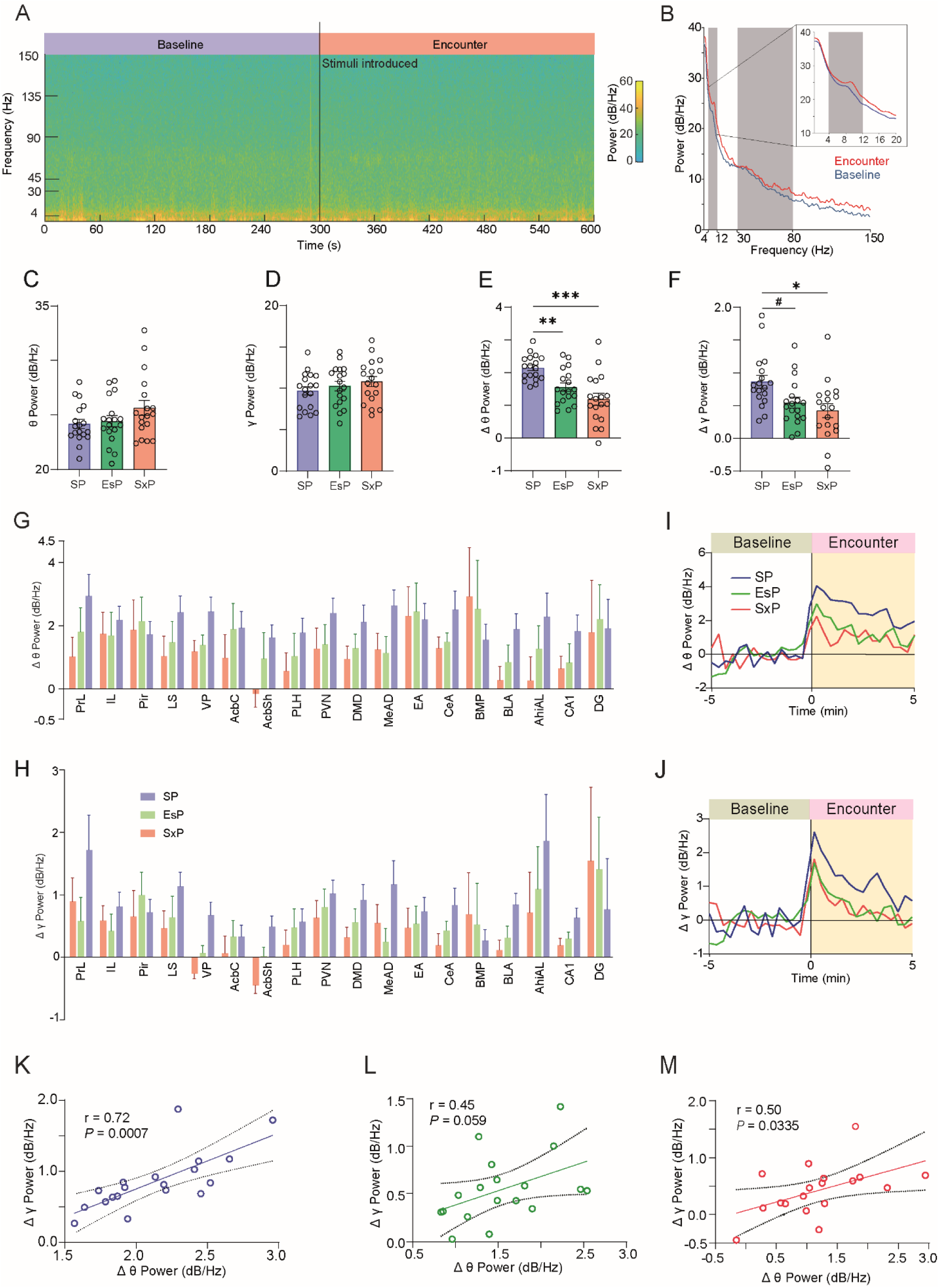
Brain region- and context-specific changes in the levels of theta and gamma power during a social encounter. A. Color-coded spectrogram of LFP signals recorded in the AcbSh during the baseline (left) and encounter (right) periods of SP task conducted by a subject. The black line at 300 s represent the time of stimuli introduction into the arena. The color-coding scale is shown on the right. B. PSD profiles of the baseline (blue) and encounter (red) periods of the example shown in A. The gray areas mark the theta and gamma ranges. The inset shows the theta range in higher resolution. C. Mean (±SEM) theta power (θP) during the baseline period for each brain region in the three contexts Kruskal-Wallis test, n = 3 tests, 87 sessions, H = 2.725, *P* = 0.2561. D. As in C, for gamma power (γP; Welch’s ANOVA, W (DFn, DFd) = 0.9496(2,33.84), *P* = 0.2561) E. Mean (±SEM) ΔθP, averaged across all brain regions, for each task. W (DFn, DFd) = 14.67 (2, 31.80), *p* <0.0001. Dunnett’s T3 multiple comparisons test, SP vs. EsP, *p* = 0.0018; SP vs. SxP, *p* = 0.0002; EsP vs. SxP, *P* = 0.2653. F. As in G, for ΔγP. W (DFn, DFd) = 5.134 (2, 33.65), *p* = 0.0113. SP vs. EsP, *p* = 0.0531; SP vs. SxP, *p* = 0.0127; EsP vs. SxP, *p* = 0.7467. G. Mean (±SEM) change in theta power (ΔθP) during the encounter period, relative to the baseline period for each brain region in the three contexts (2-way ANOVA. Contexts: F (2, 659) = 3.838, *p* = 0.0220; Brain regions: F (17, 659) = 1.727, *p* = 0.0341; Interaction: F (34, 659) = 0.4548, *p* = 0.9970) H. As in E, for change in gamma power (ΔγP; 2-way ANOVA. Contexts: F (2, 659) = 1.459, *P* = 0.2333; Brain regions: F (17, 659) = 2.050, *P* = 0.0076; Interaction: F (34, 659) = 0.5732, *P* = 0.9764). I. Super-imposed traces of ΔθP averaged across all brain regions for the SP (blue), EsP (green) and SxP (red) tasks. Time 0 min represents the time of stimuli insertion. J. As in I, for ΔγP. K. Mean ΔγP as a function of mean ΔθP during the SP task, for each brain region. Pearson’s correlation coefficient (r) and significance (*p*) are given. L. As in K, for the EsP task. M. As in K, for the SxP task. #*p*= 0.053, \**p*<0.05, \*\**p*<0.01, \*\*\**p*<0.001. See also Figs. S1, S2.

To examine the temporal dynamics of LFP rhythmicity during the various tasks, we plotted ΔθP and ΔγP as a function of time for each task and brain region. In accordance with our previous study in rats [32], we found that both ΔθP and ΔγP began to rise several seconds before stimulus introduction, peaked within 20 s from this point and gradually declined in all brain regions and tasks (Fig. S2). Thus, the dynamics of LFP rhythmicity across the session were similar among the various tasks and did not seem to reflect the behavioral dynamics (shown in Fig. 1D, H, L). We also found no significant correlation (Pearson’s, *p*>0.05) between the mean power change and speed of the subject during any task for either ΔθP or ΔγP (Fig. S3A-F).

Overall, these results suggest that theta and gamma rhythmicity during the encounter period are driven by an internal brain state that shows similar temporal dynamics across tasks, independent of the behavioral dynamics.

### LFP power changes during stimulus investigation are differentially modulated across brain regions and tasks

Despite the uniform dynamics of LFP rhythmicity in the social brain during the encounter period, it may be differentially modulated during specific behavioral events, such as stimulus investigation. We thus examined the possibility that during investigation bouts, ΔθP and ΔγP (henceforth termed ^Δ^θP and ^Δ^γP) differ between the various stimuli and tasks. As exemplified by signals recorded from the amygdalo-hippocampal area (AhiAl) shown in Fig. 3A-F, a Z-score analysis revealed elevation in theta power during investigation bouts towards social but not the object stimuli in the SP task, during investigation of both stimulus types in the EsP task and during investigation of female but not male stimuli in the SxP task. Very similar results were obtained for gamma power (Fig. S4A-F). This analysis thus suggests a bias in the response towards specific stimuli, in a task-specific manner. Interestingly, even though the same type of stimulus (a group-housed male) was used in all tasks, this stimulus elicited a clear elevation in LFP rhythmicity only during the SP and EsP tasks, but not during the SxP task. These results suggest that at least for the AhiAl, the change in LFP power was not dictated by either the stimulus type nor by its valence. To explore the stimulus-specific bias in LFP power change during each task, we calculated the difference in ^Δ^θP and ^Δ^γP between the two stimuli, separately for each brain region. Since a possible bias of LFP rhythmicity of a given brain region may be associated with a behavioral bias towards a specific stimulus, we examined the correlation between the two variables. To this end, we correlated the ^Δ^θP bias (preferred minus less-preferred) to the RDI values of each task. We found a negative correlation in a specific set of brain regions (i.e, extended amygdala (EA) and lateral septum (LS),) only for the SP task. In contrast, a positive correlation was found in a distinct set of brain regions (nucleus accumbens shell (AcbSh), AhiAl, ventral pallidum (VP) and dorsomedial hypothalamic nucleus (DMD)) during the EsP and SxP tasks. Specifically, the VP exhibited a very strong and highly significant linear correlation with RDI values during both the EsP and SxP tasks (Fig 3G). These results suggest a link between stimulus-specific bias in ^Δ^θP and behavioral preference in a task-specific manner.

**Figure 3.**
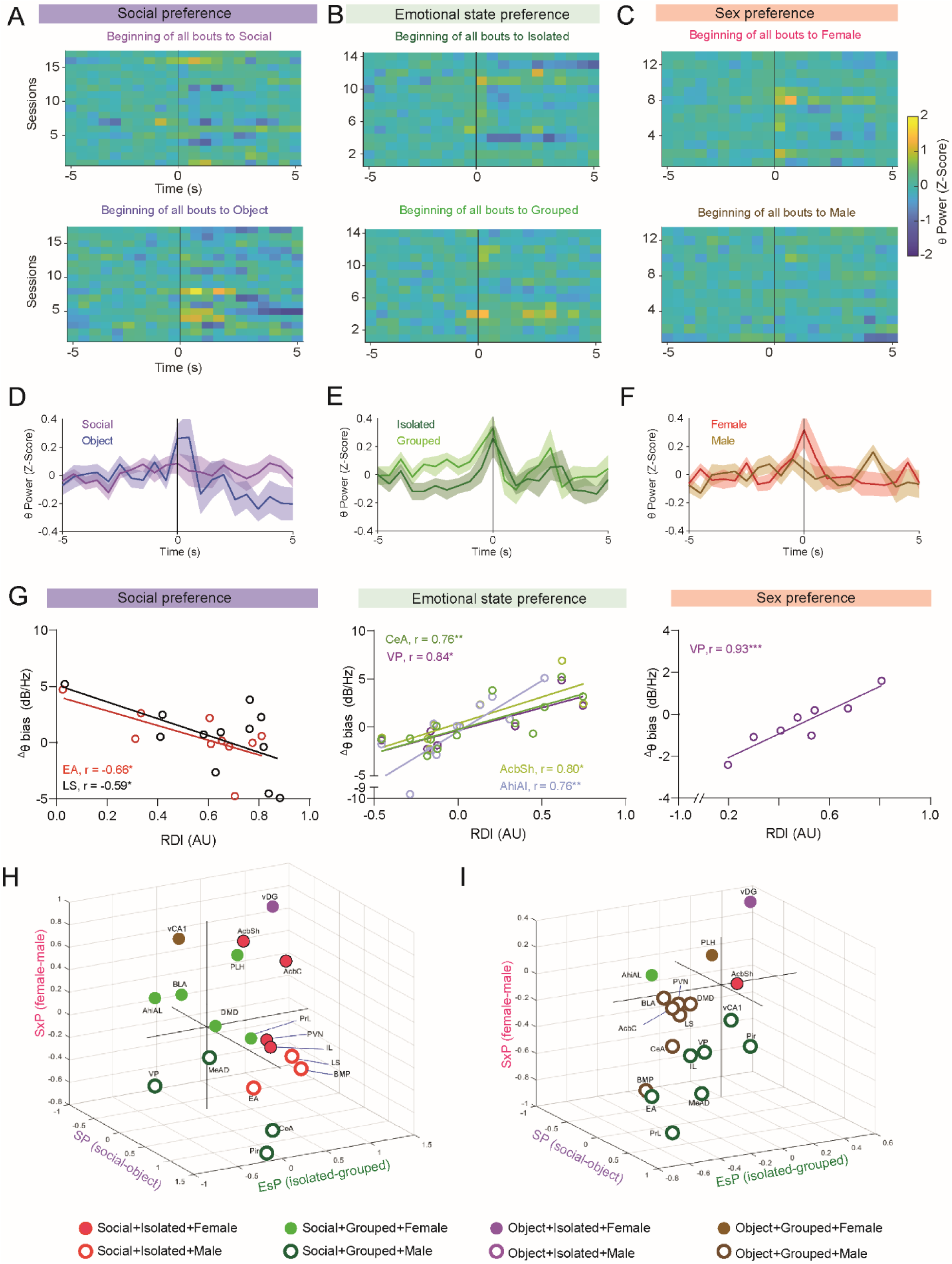
Stimulus- and task-specific LFP power changes during investigation bouts. A. Heat maps of average theta power in AhiAl, before and during social investigation bouts made by AhiAl-Implanted subjects with social (above) and object (below) stimuli during SP task sessions (n=17 sessions). Each row represents the mean Z-score of all bouts in a single session (using 0.5 s bins). Time ’0’ represents the beginning of the bout. The color code scale is on the right. B. As in A, for investigation bouts of isolated (above) and grouped (below) social stimuli during EsP task sessions (n=14 sessions). C. As in A, for investigation bouts of female (above) and male (below) social stimuli during SxP task sessions (n=13 sessions). D. Mean (±SEM) Z-score trace of the data shown in A for both stimuli. E. As in D. for the data shown in B. F. As in D, for the data shown in C. G. Correlation between mean change in theta power during investigation bouts (^Δ^θP) in specific brain regions and RDI values during the various tasks. Only statistically significant linear correlations are shown. Note that one outlier SP session which made the correlation even stronger was excluded (Fig. S4G) H. A 3D plot of the mean difference between preferred and less-preferred stimuli in ^Δ^θP. Each circle represents a given brain region, color- and shape-coded according to the combined bias across all tasks. See legend of the color and shape code of the distinct combinations below. I. As in B, for ^Δ^γP. See also Fig. S4.

To further explore this link, we plotted ^Δ^θP bias across all tasks on a 3D plot, separately for each brain region. We found that almost no region showed bias towards the object stimulus in the SP task, with the various regions being equally distributed between the two stimuli in the EsP and SxP tasks (Fig. 4B). In contrast, when ^Δ^γP was analyzed (Fig. 4C), we observed an opposite picture. Here, almost all brain regions exhibited stronger responses to the grouped and male stimuli in the EsP and SxP tasks, yet were rather equally distributed between the two stimuli in the SP task. Thus, for ^Δ^γP, most brain regions (14/18) were equally divided between those biased towards less-preferred stimuli (i.e., object+grouped+male) and those biased towards the type of stimulus used in all three tasks (i.e, social+grouped+male). The probability of such an arrangement to occur by chance is smaller than 0.001 (1-binomial test) for each of these two groups. The results thus suggest that a bias in gamma power is mostly associated with the characteristics (i.e, valence or type) of the stimulus. They also generally demonstrate opposite stimulus-dependent bias patterns between the theta and gamma rhythms during stimulus investigation, in contrast to their significant correlation when measured during the entire encounter period (Fig. 2E-G). This implies the existence of an independent active state in the social brain during stimulus investigation. Notably, of all brain regions considered, the vDG stood out as the only region biased to the combination of object/isolated/female stimuli. Moreover, this region showed an strong bias for all these stimuli in both ^Δ^θP and ^Δ^γP (Fig. 3H-I). These results suggest a unique position for the vDG in the social brain, as also supported by results presented below.

**Figure 4.**
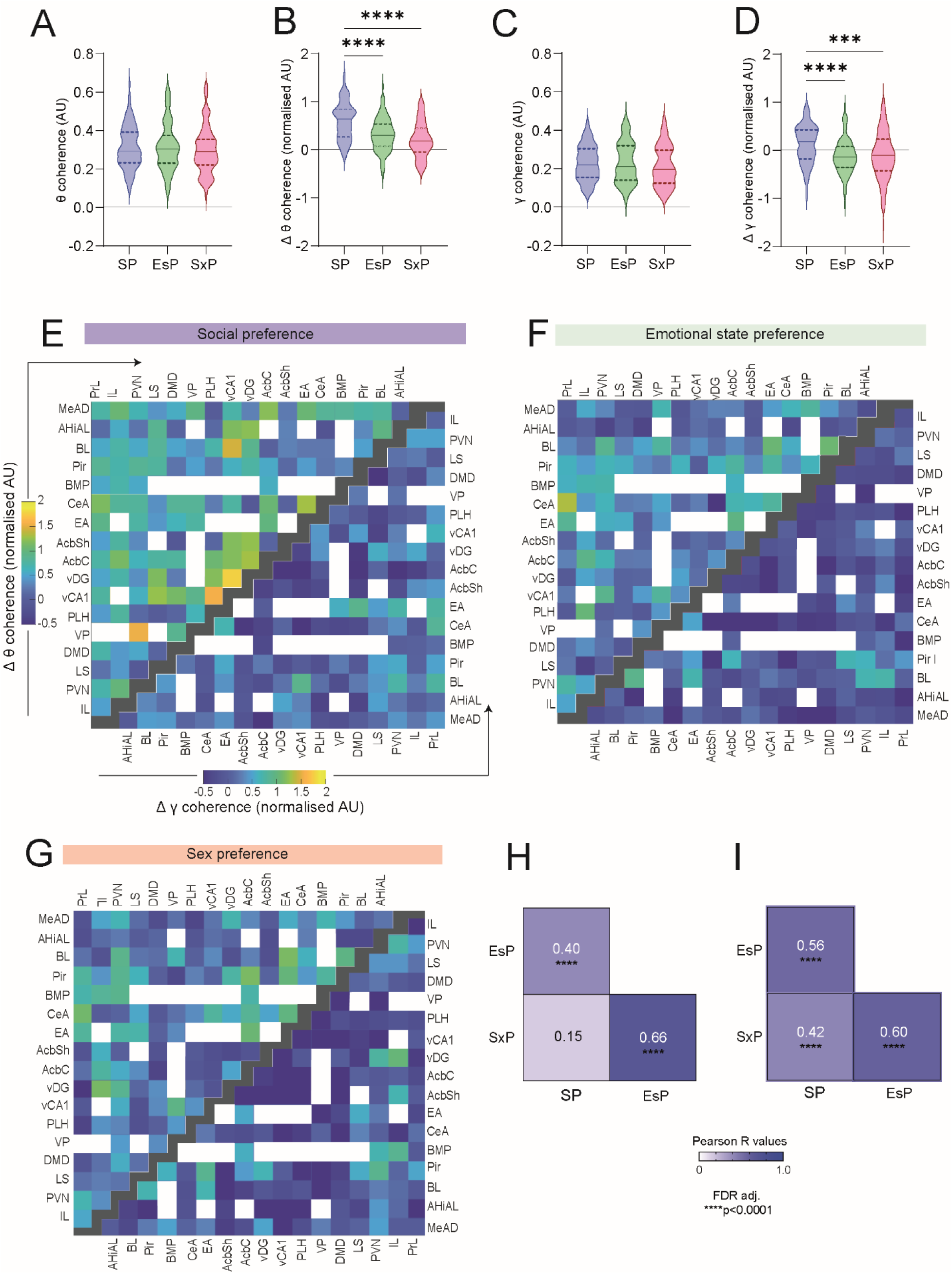
Social encounters modulate the coherence between brain regions in a social context-dependent manner. A. Mean theta coherence during the baseline period for each task, across all (n=99) pairs of brain regions recorded during all tasks (Kruskal-Wallis test, H = 0.75, *P* = 0.687). B. As in A, for mean normalized change in theta coherence (ΔθCo) during the encounter period. Note the significant difference between the SP and other tasks (****p<0.0001, Dunn’s *post-hoc* test following the main effect in a Kruskal-Wallis test). C-D. As in A and B, for gamma coherence (***p<0.001, ****p<0.0001). E. Color-coded matrix of the mean normalized ΔθCo (upper left) and ΔγCo (lower right) values for all pairs of brain regions in the SP task. Empty spots represent brain region pairs with fewer than five recorded sessions. Black spots separate between the ΔθCo and ΔγCo matrices. F. As in E, for EsP. G. As in E, for SxP. H. Coefficients and significance of Pearson’s correlations of ΔθCo across all coupled brain regions for each pair of tasks (****p<0.0001, FDR adjusted). I. As in H, for ΔγCo. See also Fig. S5.

### Social encounters modulate coherence between brain regions in a social context-dependent manner

Synchronous activity (coherence) enhances effective communication between neuronal groups in different brain regions and dynamically binds them into functional networks [38]. We, therefore, examined the coherence of LFP rhythmicity between each pair of brain regions recorded by us, in both the theta and gamma bands. During the baseline period, the mean theta coherence (θCo) between all pairs of brain regions (99 pairs with ≥5 sessions from at least two subjects in all three tasks, see Table S2)) was similar across all tasks (Fig. 4A). Thus, the subjects displayed similar global brain synchronization while exploring the arena without stimuli, in all tasks. However, the change in theta coherence (ΔθCo) during the encounter period differed significantly between tasks. While almost all pairs of brain regions exhibited increased θCo during the SP task, we observed significantly milder increases during EsP and SxP tasks, with many paired regions showing reduced θCo (Fig. 4B, E-G). Similar relationships among tasks were observed for changes in gamma coherence (γCo), although here the general tendency was one of decreased coherence during the encounter period (Fig. 4C-G). Notably, there was almost no correlation between the baseline period coherence and the change in coherence during the encounter period for any task (apart a weak correlation for SP gamma coherence; r=-0.21, p=0.0111; not shown). This suggests that the encounter-induced coherence change represents an internal state, independent of the resting state. Finally, when calculating the correlations in ΔθCo across all paired regions between the various tasks, we found a statistically significant high correlation between SxP and EsP, while no correlation was found between SP and SxP. A milder but significant correlation was found between SP and EsP (Fig. 4H). In contrast, all correlations were found significant for ΔγCo. These results suggest a gradual shift from SP to EsP and to SxP when in ΔθCo brain pattern is measured across the whole encounter.

We, therefore, examined the encounter-induced coherence changes for each brain region separately, by comparing the change in coherence between a given region and all other regions across tasks in the theta (Fig. S5A) and gamma (Fig. S5B) bands. We found that differences between tasks were brain-region specific, with most (13/18) regions showing significant differences (after FDR corrections) between at least two tasks in ΔθCo and one third (6/18) showing differences in ΔγCo. Notably, in all cases, we found significantly higher coherence changes during the SP task, as compared to at least one task, and in many cases, to both other tasks. Thus, a subset of the recorded brain regions displayed differential changes in theta or gamma coherence among the social contexts, with this change being majorly increased in the SP task, as compared to the other two tasks.

### Brain-wide coherence changes during investigation bouts reflect the social context

To explore possible modulation of LFP coherence during investigation bouts, we calculated the mean ^Δ^θCo between each pair of brain regions during all investigation bouts towards a given stimulus, similarly to how we analyzed the power changes (Fig. 3). Since data had been collected for a large number of brain-region pairs (99 pairs), we focused on pairs showing a mean coherence change that crossed a cutoff value ± 1.5*standard deviation (SD) for each stimulus (about 20% of the pairs). When plotting the bias in ^Δ^θCo and ^Δ^γCo between the two stimuli in each task on a 3D plot (Fig. S6A-B), we observed a much wider distribution than was found for power (Fig. 3H-I), suggesting distinct principles of distribution. To further explore this possibility, we examined all pairs of brain region that passed the aforementioned threshold, separately for each stimulus (Fig. 5A). Surprisingly, the three stimuli of the same type (i.e, social, grouped, male) did not share even a single pair of brain regions that passed the ^Δ^θCo cutoff value. Similarly, the preferred stimuli (i.e, social, isolated, female) also did not share even a single pair among them. In contrast, multiple pairs of brain regions shared similar changes in theta coherence between both stimuli used in each task (Fig. 5A). For example, CeA-PrL and MeAD-VP showed increased coherence for both social and object stimuli, BLA-LS and EA-Acbc showed increased coherence for both isolated and grouped stimuli and Pir-AcbC and EA-AcbC exhibited the strongest increase in theta coherence for both male and female stimuli. Similar results were observed for ^Δ^γCo (Fig. S6C). Thus, changes in coherence during stimulus investigation seem to be dictated by the social context rather than by stimulus characteristics, such as its type or valence.

**Figure 5.**
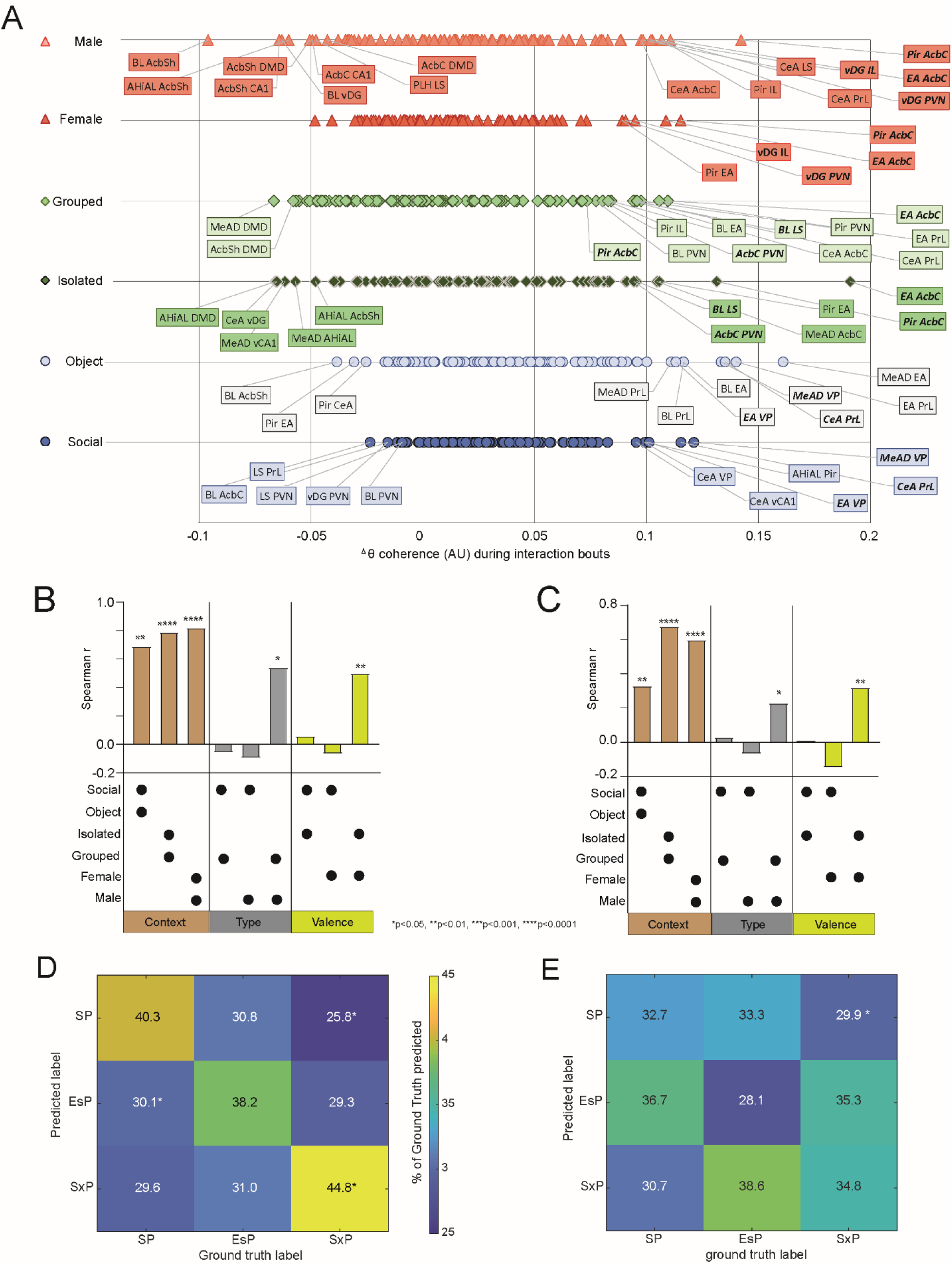
Coherence changes during social investigation are informative regarding the social context. A. Distribution of changes in theta coherence during investigation bouts (^Δ^θCo) between each pair of brain regions, plotted separately for each stimulus used in the SP (blue), EsP (green) and SxP (red) tasks. The names of brain region pairs which passed the mean cutoff value ±1.5*SD are labeled, with those showing similarly high ^Δ^θCo values for both stimuli of the same task in bold. B. Spearman’s correlation coefficients of mean ^Δ^θCo across all paired brain regions, for couples of stimuli which were either used in the same task (left, brown bars), of the same type (middle, gray) or having the same valence (right, yellow). The correlated two stimuli are denoted by asterisk below each bar, while the statistical significance of the correlation is marked above the bars. C. As in B, for ^Δ^γCo. D. A color-coded confusion matrix for a multi-class Random forest classifier employed for predicting the social context from ^Δ^θCo values across all brain regions and stimuli. The scale of the accuracy’s color code is shown to the right. The percentage of cases a label was predicted for each ground truth are marked in the middle of each spot. *p<0.05, Mann-Whitney test, FDR adjusted. E. As in D, for ^Δ^γCo. See also Figs. S6, S7.

For quantitative examination of this possibility, we calculated the correlation across all brain regions for either ^Δ^θCo (Fig. 5B) and ^Δ^γCo (Fig. 5C), between pairs of stimuli which share common context, type or valence. We found strong and highly significant correlations between all pairs of stimuli used in the same task (sharing context). In contrast, among the three stimuli of the same type, only grouped (ESP) and male (SxP) showed significant correlation. Similarly, among preferred stimuli, only isolated (EsP) and female (SxP) showed significant correlations. Notably, in both of these cases the correlation was weaker than the correlation between any pair of stimuli sharing the same context (Fig. 5B). Similar results were found for ^Δ^γCo (Fig. 5C). Thus, coherence changes during stimulus investigation in both bands had the strongest association with the context of the social interaction, relative to any characteristic of the stimulus.

Finally, we employed a Decision trees (multi-class Random forest) classifier to examine if ^Δ^θCo and ^Δ^γCo contain information which may be used to discriminate between the various contexts or stimuli. First, we validated that the model achieved good (∼60%) and significant accuracy in predicting the social stimulus vs. object in the SP task using either ^Δ^θCo or ^Δ^γCo. Notably, the classification of the object vs. social was not accurate, suggesting that the presence of the social stimulus (social context) mask the object classification (Fig. S7A-B). Then, we used the same model for predicting the social context (SP, EsP and SxP) and found that using ^Δ^θCo (Fig. 5D), but not ^Δ^γCo (Fig. 5E), allowed the model to predict the right context better than any other context, and that this prediction was the only one achieving more than a chance level (33.3%) accuracy (although only SxP classification was statistically significant). In contrast, the same model worked poorly for predicting stimulus identity among all six stimuli (Fig. S7C-D). Using the LFP theta power (^Δ^θP) for predicting the social context by the same model achieved good and significant classification only for the SP (Fig. S7E), while using both theta power and coherence allowed accurate prediction of both SP and SxP contexts (Fig. S7F). These results suggest that LFP rhythmicity in the theta range, especially the coherence between the various brain regions, is informative regarding the social context of the animal more than regarding the identity or valence of the social stimuli.

### Analysis of Granger causality suggests that specific brain regions serve as hubs

The coherent LFP rhythmicity in the social brain can be dominated by specific regions serving as hubs, thereby preceding other regions in terms of rhythmic neural activity. To identify hub candidate regions, we first selected brain regions which are statistically over-represented (see Methods) among pairs of regions exhibiting strong (mean ± 1.5*SD) bias in any task, separately for ^Δ^θCo (Fig. 6A) and ^Δ^γCo (Fig. 6B). We then examined the dependence of LFP rhythmicity of each of these regions in terms of preceding rhythmicity of other regions, by calculating the change in Granger causality (GC) [43] during the encounter period (Fig. 6C-E), as compared to the baseline. We found distinct patterns of statistically significant changes in GC (encounter vs. baseline periods) between the various tasks (Fig.6-H). Some regions, however, presented significant GC changes in all tasks, suggesting that they might function as hubs. For example, the vDG and AcbC participated in significant GC changes in both theta and gamma rhythms in all tasks. At the same time, PrL and AhiAl were explicitly involved in theta GC changes in all tasks. Interestingly, theta GC changes from vDG to AhiAl increased during a SP task but decreased during a EsP task, while theta GC changes from PrL to vDG decreased during both EsP and SxP tasks. These results suggest that these brain regions dictate LFP rhythmicity in the social brain during social investigation.

**Figure 6.**
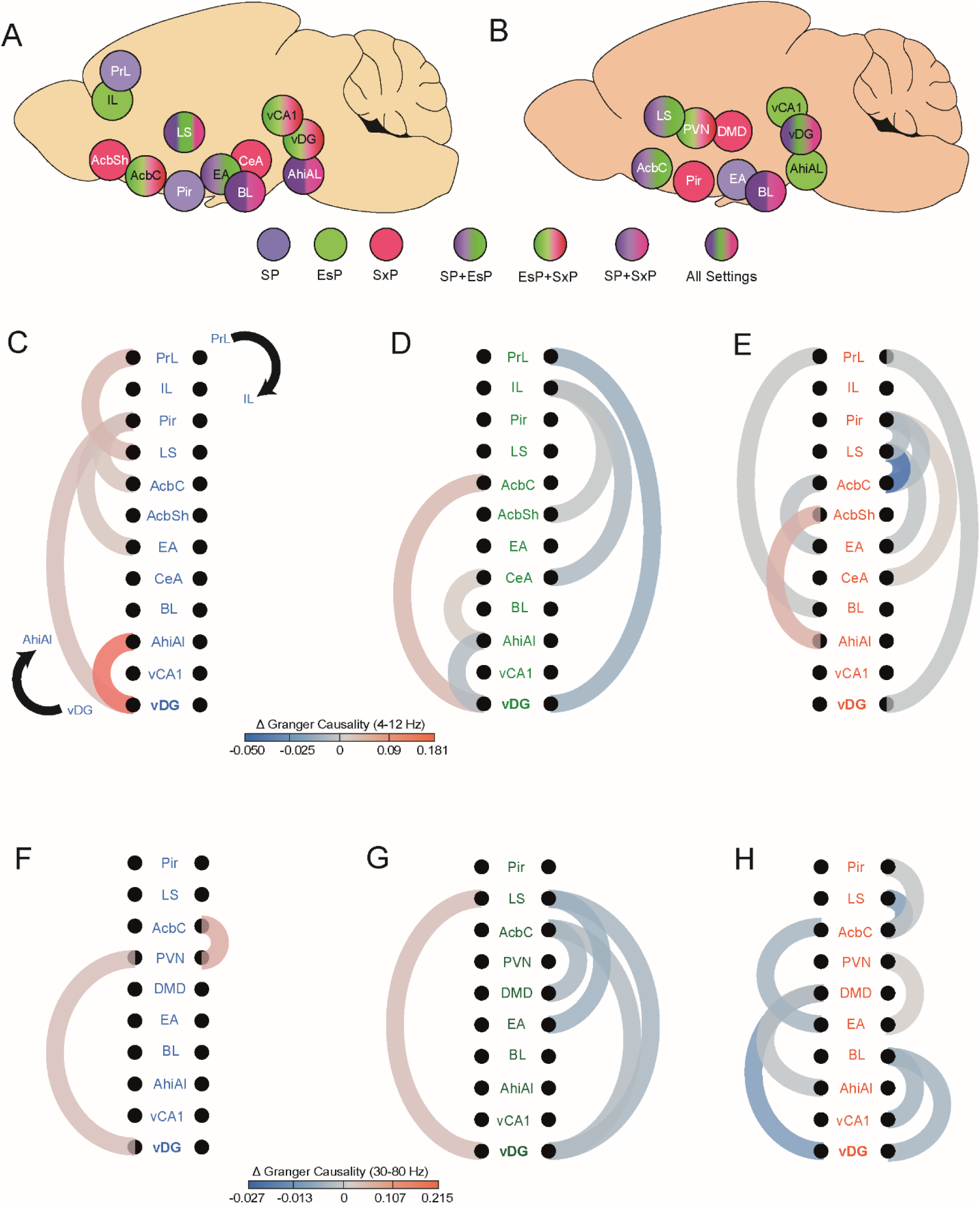
Distinct patterns of changes in Granger causality (GC) during the encounter period among tasks. A. Schematic representation of the brain regions over-represented among the pairs that exhibited strong (mean ± 1.5*SD) theta coherence bias towards one of the stimuli in any task. The regions are color-coded according to the task in which they were over-represented. B. As in A, for gamma coherence. C. Schematic representation of significant changes (encounter vs. baseline) in theta band GC during a SP task, among the regions listed in A. The direction of the GC changes in shown by a black arrow (top to bottom on the right and bottom to top on the left). D. As in C, for EsP. E. AS in C, for SxP. F-H. As in C-E, for gamma band See also Fig. S8

To further explore this possibility, we have calculated the difference in GC change during encounter between the two directions (from area 1 to area 2 and vice versa), for all couples of brain regions across all tasks and rhythms (Fig. S8A-C). After applying FDR correction for multiple comparisons, we found only vDG to LS, for gamma rhythmicity of the EsP task, which was significantly higher in the vDG-LS direction than in the opposite direction (Fig. S8D).

### Context-specific synchronization of LFP rhythmicity in the ventral dentate gyrus with precise behavioral events

To further examine the candidate hub regions, we exploited our ability to determine the exact timing of each investigation bout to quantify the synchronized modulation of LFP rhythmicity, relative to these events. Thus, we compared the modulation of theta and gamma power in all regions associated with significant GC changes (Fig. 6) relative to a defined battery of specific behavioral events (Fig. 7A). These events included the beginning and end of investigation bouts towards specific stimuli, as well as transitions between stimuli. We found a main effect in ANOVA for multiple events, although in most cases, none of the regions showed significance *in post-hoc* analysis (see Table S3). One region, the vDG, did, however, exhibit significant differences between stimuli. The vDG displayed significantly higher theta and gamma powers at the end of investigation bouts of social stimuli, as compared to object stimuli, specifically in the SP task (Fig. 7B-G and S7A-B). The same region also exhibited decreased theta and gamma powers at the beginning of transitions from isolated to grouped stimuli, as compared to non-transitional bouts, specifically in the EsP task (Fig. 7H-M and S7C-D).

**Figure 7.**
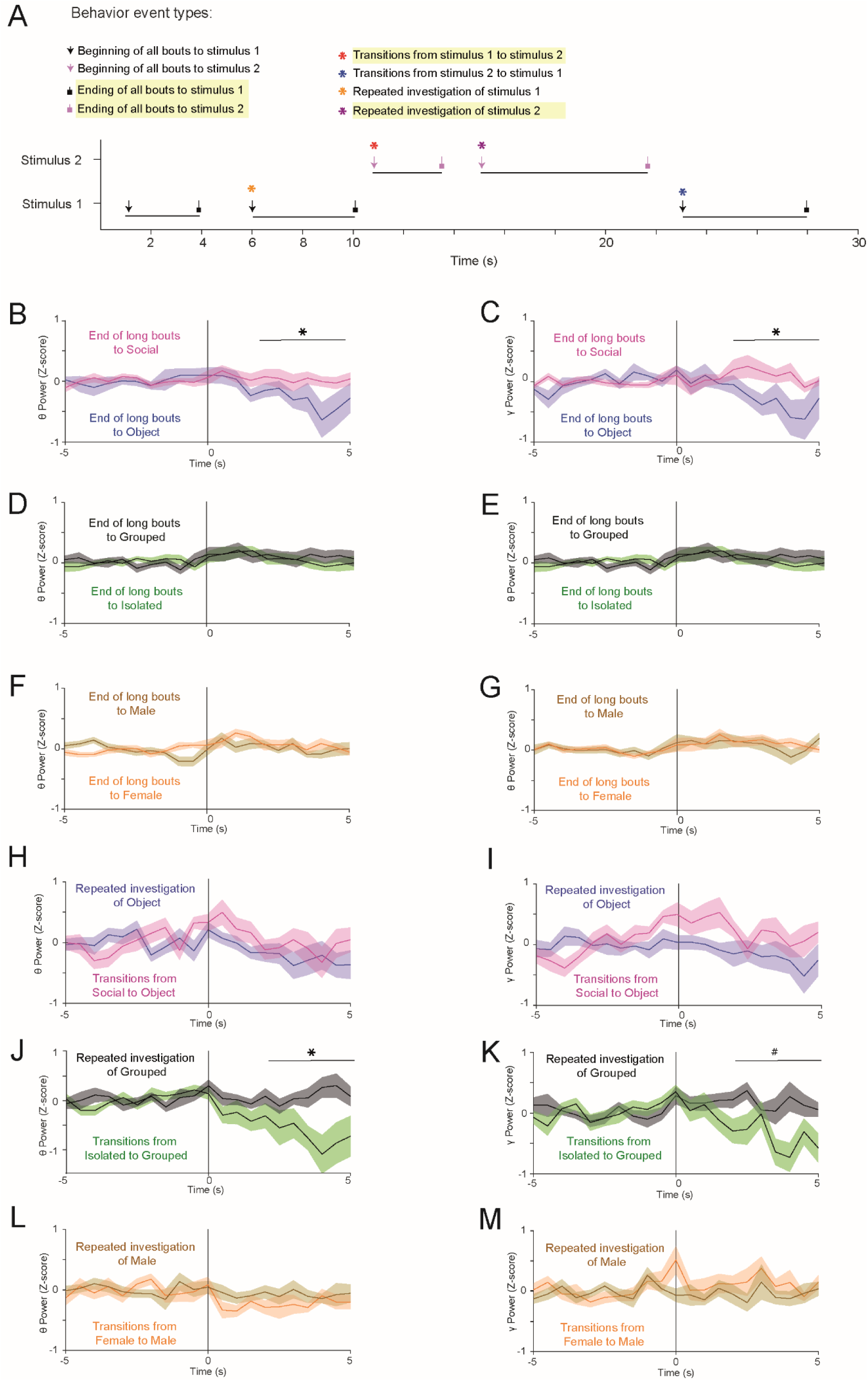
Context-specific differences in vDG LFP rhythmicity in specific behavioral events. A. Color-coded scheme of specific behavior event types, with **↓** indicating the beginning of an, while **Ʇ** indicating the end of an investigation bout, and **\*** indicating the beginning of a bout after transition between stimuli or repeated investigation of the same stimulus. Events showing significant differences in vDG LFP power are highlighted in yellow. B. Super-imposed traces of the mean (±SEM) Z-score of changes in vDG theta power at the end of long bouts towards either a social stimulus (pink) or an object stimulus (purple) in the SP task. Time 0 represents the end of the bout, while the 5 s period before time 0 was considered as baseline. \**p*<0.05, Student’s t-test between the mean Z-score values averaged over the last 3 s of the traces. C. As in B, for gamma power in the vDG during SP tasks. D-E. As in B-C, for the EsP task. F-G. as in B-C, for the SxP task. H-I. As in B-C, for changes in LFP power at the beginning of repeated vs. transitional (between stimuli) investigation bouts of social and object stimuli across SP task sessions. J-K. As in H-I, for the EsP task. \*\**p*<0.01 by a Mann-Whitney test following the main effect in ANOVA. L-M. As in H-I, for the SxP task. See also Fig. S9.

These results, together with those shown in Fig. 3B-C and Fig. 6, suggest that the vDG may function as a hub in the social brain network by coordinating rhythmic neural activity of the network in a social context-dependent manner.

## Discussion

In this study, we used multi-site electrophysiological recordings from the murine social brain to seek system-level neural correlates of three distinct aspects of social interaction, namely, the type of the social stimulus, its relative valence (preference) and the social context. To distinguish between these three aspects, we relied on three social discrimination tasks (i.e., SP, EsP, and SxP) in which male mice clearly prefer one of two distinct stimuli. This design enabled us to employ the same type of social stimulus, a novel group-housed male mouse, in all three tasks, with this stimulus being the preferred stimulus in the SP task and the less-preferred stimulus in the other two tasks. Importantly, all three tasks took place in the same experimental arena, which enables uniform interactions between the subject and the stimuli, i.e., stimulus investigation by the subject [42]. Consequently, as much as we could measure, subject behavior was almost identical in all three tasks. Therefore, behavioral differences cannot explain the significant differences in the patterns of rhythmic LFP signals observed among the different tasks.

We analyzed LFP signals at three different time resolutions, specifically, across the whole session, during stimulus investigation, and during specific behavioral events, such as at the beginning and end of investigation bouts. When analyzing the power of both theta and gamma rhythms over an entire session, some aspects seemed to be dictated by a general internal state. In accordance with previous studies by us and others [31, 32], virtually all brain regions exhibited higher level of theta and gamma power during the encounter period, as compared to the baseline period. Our observation that the level of enhanced power was both brain region- and task-specific strongly suggests that the power elevation was not caused by enhanced electrical noise or any other artifact but rather by a genuine internal state of the animal. The uniform dynamics of both theta and gamma power changes across all brain regions and tasks during the encounter period further supports the existence of a general internal state which is independent of behavioral dynamics or context. Notably, we observed significant correlations across brain regions between theta and gamma power changes in all tasks, suggesting that both rhythms are similarly influenced by the internal state. In agreement with our previous studies in both rats and mice [30, 32], theta and gamma power maintained their high levels for a time, even after removal of the stimuli from the arena (not shown), further supporting an encounter-induced general internal state, which slowly fades away. This state did not seem to be caused by subject movement, as we found no correlation between subject speed and changes in theta or gamma power for any brain region.

While the dynamics of the internal state seemed to be similar across the distinct contexts, other aspects of the general (session-wise) changes in theta and gamma power exhibited context-specific characteristics. For example, the general changes in both power and coherence were highest in the SP task, suggesting a higher level of the internal state. Assuming that the general state reflects social motivation, these results are somewhat surprising, given how the SP task involved only one social stimulus and reasoning that among the various stimuli tested, the female would be the most attractive to the male subjects. Our interpretation is that the SP task is simpler in terms of social motivation, as it requires the animal to choose between an inanimate object and a conspecific, while the other two tasks involve two social stimuli, thereby presenting the subject with a more challenging dilemma. The higher confidence of the subject during the SP task is in accord with the simpler pattern of theta coherence changes observed during this task (seen as a general increase across almost all brain region pairs). Overall, these results suggest that the internal state level may distinguish between some contexts, which is in accordance with the ability of the Random forest model to predict only the SP context based on the arousal-induced LFP power. Nevertheless, the changes in theta and gamma power across the encounter period did not differ between the EsP and SxP tasks, and thus cannot be the sole basis for the context-specific responses to social cues.

Notably, a recent paper [22] that employed similar recordings during the SP test, used the power, coherence and GC data (termed *Electome network*) from various regions of the social brain in a machine-learning model to discriminate between social and object investigation. In accordance with our results, this study reported that the model’s precision was higher for the social than for the object, thus suggesting that the social stimulus masks the object, which may be attributed to the context effect.

Analysis of the power change, specifically during stimulus investigation, yielded a different picture than did session-wide analysis. First, we found no correlation between theta and gamma power changes during these periods, suggesting a distinct state of active sensing which characterizes stimulus investigation. Moreover, although both theta and gamma power changes across brain regions showed bias to specific combinations of stimuli, they did so in distinct manners. While theta power was biased towards the preferred stimulus in the SP task, with almost no region (other than hippocampal areas) showing a higher level during investigation of object stimuli, the gamma power was clearly biased towards the less-preferred stimuli in the EsP and SxP tasks (grouped and male stimuli), while showing a mixed preference between stimuli in the SP task. Thus, as related to gamma power, the social brain may be divided between regions associated with the valence of stimulus (biased towards less-preferred stimuli) and brain regions associated with the type of stimulus. It should be noted that theta rhythmicity is thought to reflect top-down processes, such as arousal and attention, which are regulated by brain wide-active neuromodulators and recruit distributed brain networks [48–52]. In contrast, gamma rhythmicity is considered a bottom-up process [53, 54], associated with the synchrony of local inhibitory networks [53–56]. This distinction may explain why theta and gamma rhythms reflect stimulus characteristics in an opposing manner.

In accordance with our hypothesis, that coherent theta and gamma rhythms couple various regions of the social brain in a social context-dependent manner, we found that the correlation in the coherence change during stimulus investigation was strongest between the two stimuli in each task, even the EsP and SxP tasks. In contrast, there were weaker correlations, if any, among the three stimuli of the same type (social, grouped, male) or the preferred stimuli (social, isolated, female). The fact that the same correlation pattern was observed for the coherence of both theta and gamma rhythms supports the validity and significance of the observation. Moreover, using a Decision Tree classifier, we demonstrated that the theta coherence between the recorded areas could generate predictions regarding the social context, but not the specific stimulus, which are accurate above the chance level. The limited accuracy of the model may be attributed to the restricted number of recorded regions. Thus, we expect that a more comprehensive analysis of the coherence within the social brain will be able to generate highly accurate prediction of the social context. Moreover, GC analysis, representing causal time relationships between various brain regions, also suggests distinct patterns of changes across the various contexts. Altogether, these results are in accordance with the idea that the social brain processes information during stimulus investigation in a context-dependent manner dictated by the context-dependent pattern of coherence within the network. Such a mechanism may explain how the same stimulus induces distinct patterns of brain activity in different social contexts, which then elicits distinct behavioral responses to a stimulus.

Finally, the coherence changes and GC analyses led us to identify a small subset of brain regions that seem highly influential within the network during the various tasks. Of these, the vDG and AcbC were involved in significant GC changes during all tasks in both the theta and gamma bands, and thus may serve as hubs that influence the activities of other regions. Analysis of LFP power in relation to a battery of specific behavioral events demonstrated that while the small group of brain regions considered showed differential responses as a whole, the vDG was the only region that alone showed statistically significant responses. Together with its strong bias towards specific stimuli, as demonstrated for both theta and gamma power during investigation bouts (Fig. 4B-C), these results suggest a role for the vDG in orchestrating neural activity across the social brain during social behavior. This conclusion agrees with previous studies reporting a central role of the dentate gyrus in social behavior [57–59], and specifically in social discrimination [60, 61]. Notably, multiple studies have implicated the DG in coding contextual changes. For example, DG neurons were shown to rapidly detect and encode contextual changes [62], while knocking out NMDA receptors specifically in DG granule cells abolished the ability of mice to distinguish between two similar contexts [40]. Moreover, hypothalamic supramammillary neurons projecting to the DG were shown to be activated by contextual novelty [39], while the activity of ventral hippocampal neurons was shown to process information in a social context-sensitive manner [63]. These studies are thus in line with our findings regarding the involvement of the vDG in context-dependent social behavior.

In conclusion, our results suggest that the distribution of LFP rhythmic activity in the social brain and, most specifically, the synchronization between the various regions is context-specific and may thus mediate context-specific processing of social information, leading to social context-dependent social responses and behavior.

## Methods

### Animals

Adult male and female CD1 mice (12-14 weeks old) were acquired from Envigo (Rehovot, Israel). All mice were housed in groups of 3-5 in a dark/light 12-hour cycle (lights on at 7 pm), with *ad libitum* food and water. Following surgery, implanted mice were housed in isolation so as to not disturb the implanted EAr. Experiments were performed in the dark phase of the dark/light cycle in a sound- and electromagnetic noise-attenuated chamber. All experiments were approved by the Institutional Animal Care and Use Committee of the University of Haifa (Ethical approval #616/19).

### Surgery

Mice were anesthetized using isoflurane (induction 3%, 0.5%-0.8% maintenance in 200mL/min of air; SomnoSuite) and placed over a custom-made heating pad (37°C) in a standard stereotaxic device (Kopf Instruments, Tujunga, CA). Two burr holes were drilled for placing the ground and reference wires (silver wire, 127 µm, 300-500 Ω; A-M Systems, Carlsborg, WA). Two watch screws (0-80, 1/16”, M1.4) were inserted into the temporal bone. The coordinates for Prl (AP= 2mm, ML= -0.3, DV= -1.8), AcbC (AP= 1, ML= -2.3, DV= -4.7), Pir (AP= -2, ML= -3.3, DV = - 5) and CA1 (AP= -3, ML= -3.3, DV = -4.7) were indicated over the left hemisphere using a marker. The skull covering these marked coordinates was removed using a dental drill, and the exposed brain was kept moist with cold, sterile saline. We custom-designed the EA [41] from 16 individual 50 µm formvar-insulated tungsten wires (50-150 kΩ, #CFW2032882; California Wire Company). Before implantation, the EAr was dipped in DiI (1,1’-Dioctadecyl-3,3,3’,3’-tetramethylindocarbocyanine perchlorate; 42364, Sigma-Aldrich) to visualize electrode locations *post-mortem*. The reference and ground wires were inserted into their respective burr holes. The EAr was lowered onto the surface of the exposed brain using a motorized manipulator (MP200; Sutter instruments). The dorsoventral coordinates were marked using the depth of the electrode targeting the PVN (AP= -1 mm, ML= -0.3), which was lowered slowly to -4.7 mm. The EA and exposed skull with the screws were secured with dental cement (Enamel plus, Micerium). Mice were sub-cutaneously injected with Baytril (5mg/kg; Bayer) and Norocarp (5 mg/kg; Carprofen, Norbrook Lab) post-surgery and allowed to recover for three days.

### Electrophysiological and video recordings

Following brief exposure to isoflurane, subjects were attached to the headstage (RHD 32 ch, #C3314, Intan Technologies) through a custom-made Omnetics to Mill-Max adaptor (Mill-Max model 852-10-100-10-001000). Behavior was recorded using a monochromatic camera (30 Hz, Flea3 USB3, Flir) placed above the arena. Electrophysiological recordings were made with the RHD2000 evaluation system using an ultra-thin SPI interface cable connected to the headstage board (RHD 16ch, #C3334, Intan Technologies). Electrophysiological recordings (sampled at 20 kHz) were synchronized with recorded video using a TTL trigger pulse and by recording camera frame strobes.

### Experiment design

We recorded the behavior and neural activity of 14 males in the SP task, 13 males in the EsP task, and 11 males in the SxP task (Table S1), while targeting 18 distinct brain regions. All the stimuli used for the tasks were unfamiliar to the subject mice. In experiments, the mice were briefly exposed to isoflurane, and the EAr was connected to the evaluation system. After 10 minutes of habituation, the recordings started in the arena (30 x 22 x 35 cm) with empty triangular chambers (12 cm isosceles, 35 cm height), as previously described [42]. The triangular chambers had one face ending with metal mesh (18 mm x 6 cm; 1 cm x 1 cm holes) through which the mice interacted with the stimuli. The test was divided into two 5 min periods, a baseline period (pre-encounter) and a period of encounter with the stimuli. The stimuli for the SP task were a novel group-housed male mouse (social) and a Lego toy (object). For the ESP task, isolated (7-14 days) male and group-housed male mice served as stimuli. Finally, for the SxP task, group-housed male and female mice were used as stimuli. Each subject was evaluated for three sessions of each task. The subjects first performed SP and free interactions, with 10 min between these tasks for three sessions, and then EsP and SxP tasks were performed likewise. Each day four sessions were recorded, two in the morning and two in the afternoon, six hours apart (See Fig. S1B). The free interaction data were not used in this study. We excluded sessions from further evaluations when there was a removal of the headstage from the EA by subjects or in a case of a missing video recording from a session . This accounts for the unequal number of sessions and subjects across tasks.

### Histology

Subjects were transcardially perfused, and their brains was kept cold in 4% paraformaldehyde for 48 h. Brains were sectioned (50 µm) horizontally (VT 1200s, Leica). Electrode marks were visualized (DiI coated, Red) against DAPI-stained sections with an epifluorescence microscope (Ti2 eclipse, Nikon). The marks were used to locate the respective brain regions, based on the mouse atlas. Out of all implanted electrodes (256), 9% (23 electrodes from 14 mice) were found to be mistargeted and 36% (93) were non-functional (Table S1).

### Behavioral analysis

Subject behavior was tracked using the TrackRodent algorithm [42] for tethered mice. Further parameters of behavior, like duration of interaction, interaction bouts, distance traveled by the subjects, subject speed, transitions between stimuli, and RDI values were calculated from the tracked results with custom codes written in MATLAB 2020a.

### Electrophysiological data analysis

Only brain regions recorded for more than 5 sessions across at least 3 mice were analyzed. All signals were analyzed with codes custom-written in MATLAB 2020a. We excluded the signals recorded during 30 seconds around stimulus removal and insertion times, to avoid any effect of this action. First, the signals were down-sampled to 5 kHz and low-pass filtered to 300 Hz using a Butterworth filter. The power and time for the different frequencies were estimated using the ’spectrogram’ function in MATLAB with the following parameters: Discrete prolate spheroidal sequences (dpss) window = 2 s; overlap = 50% ; increments = 0.5 Hz; and time bins = 0.5 s. The power of each frequency band (theta: 4-12 Hz and gamma: 30-80 Hz) was averaged for both the baseline and encounter periods (5-min long each). Changes in theta (ΔθP) and gamma (ΔγP) powers for each brain region were defined as the mean difference in power between the encounter and baseline periods (Fig. 2C-D, H-I). For Z-score analysis of ^Δ^θP and ^Δ^γP during investigation bouts for a given stimulus we used the pre-bout 5 s period as baseline, and averaged the Z-score across all bouts with the same stimulus in each session. Notably, throughout the study we have analyzed only investigation bouts that were longer than 2 s, for two reasons: 1) only >2 s bouts showed statistically significant differences between the stimuli in the various tasks and 2) only >2 s bouts allow a reliable calculation of theta coherence. LFP power (^Δ^θP and ^Δ^γP) for specific bouts with each stimulus was estimated by calculating the difference between the average power per second during an investigation bout (which was longer than 2 s) during the encounter period and the average power per second for investigation of both empty chambers in the baseline period of the same session, followed by averaging these values over all sessions (Fig. 4).

### Coherence analysis

We used the ’mscohere’ function of MATLAB to estimate coherence values using Welch’s overlapped averaged periodogram method. The magnitude-squared coherence between two signals, x, and y, was defined as follows:

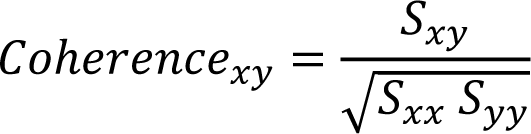

where 𝑆_𝑥𝑦_ is the cross-power spectral density of x and y, 𝑆_𝑥𝑥_ is the power spectral density of x and 𝑆_𝑦𝑦_is the power spectral density of y. All coherence analysis was quantified between brain regions pairs involved in at least five sessions of behavior tasks. Coherence for the baseline period was quantified as the average coherence of all brain region pairs for each context (Fig. 5A and F). Changes in coherence (ΔθCo and ΔγCo) during the encounter period (Fig. 5B and G) between a pair of brain regions were calculated as follows:

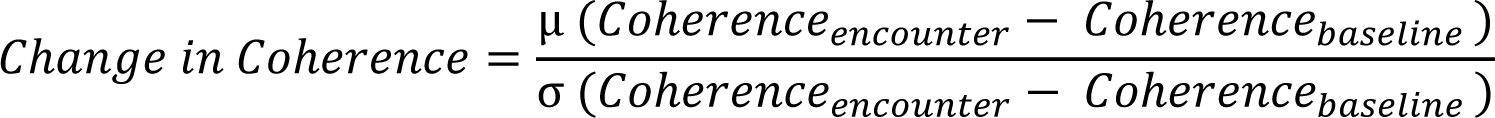

where, Coherence_encounter_ is the absolute coherence value between a pair of regions within a frequency band during whole encounter period. Coherence_baseline_ is the absolute coherence value between a pair of regions within a frequency band during an entire encounter period. The change in coherence for specific bouts (^Δ^θCo and ^Δ^γCo) to each stimulus was estimated by calculating the difference between the average coherence per second during an investigation bout (≥2 s) in the encounter period and the average coherence per second for investigation with both empty chambers during the baseline period of the same session, followed by averaging these values over all sessions (Fig. 6A and S5C). Brain regions that displayed higher frequencies of crossing the threshold of mean ± 1.5 SD ^Δ^θCo and ^Δ^γCo, based on a binomial distribution test, were considered to be hubs in the coherent social brain (Fig. 7A-B).

### Inter-regional pairwise conditional Granger causality

We employed the multi-variate GC toolbox [43] to calculate GC values separately for baseline and encounter periods between brain regions separately for each task and rhythm. To this end, we selected brain regions most represented among brain region pairs that crossed the mean ± 1.5*SD threshold for the difference in coherence change between preferred and less preferred stimuli in any task, separately for ^Δ^θCo and ^Δ^γCo. For GC analysis, LFP signals were measured at a reduced sampling rate of 500 Hz. We used the “tsdata_to_infocrit” function to determine the model order of the vector autoregressive (VAR) model. The median model order for all three tasks was 38 (Bayesian information criterion). To further fit the VAR model to our multi-session, multivariate LFP data, the “tsdata_to_var” function of LWR (Levinson-Whittle recursion) in the regression mode and a median model order of 38 was used separately for the baseline and encounter periods of each task. Next, we estimated the autocovariance sequence of the fitted VAR model with the “var_to_autocov” function. To maximize the computational efficiency of the function, an acmaxlags of 1500 was chosen. This process did not violate the autocovariance VAR model, as was estimated by the “var_info” function. Finally, we calculated the pairwise conditional frequency-domain multivariate GC matrix using the “autocov_to_spwcgc” function, and summed the GC for the relevant frequency band (theta or famma) using the “smvgc_to_mvgc” function.

### Neural responses to behavioral events

We divided all investigation bouts into specific behavioral events, such as the start and end of an investigation bout or transition from one stimulus to the other (Fig. 7A, nine distinct types). We aligned LFP power and behavior events for each stimulus by calculating mean power 5 s before and 5 s after the beginning (or end) of all investigation bouts (0.5 s bins) in a session. Furthermore, for each bout, the mean power was normalized using Z-score analysis, where a pre-bout duration of 5 s served as baseline (Table S2).

### Statistical analysis

Statistical analysis was performed using GraphPad Prism 9.5. To test for normal distribution of the data, we used the Kolmogorov-Smirnov and Shapiro-Wilk tests. Table S3 summarizes the specific tests conducted for each figure. A paired t-test or Wilcoxon matched-pairs signed rank test was used to compare different stimuli or conditions for the same group. An unpaired t-test or Mann-Whitney test was performed to compare a parameter between distinct groups. For comparison among multiple groups and parameters, ANOVA (normal distribution), Welch’s ANOVA (assuming unequal variance), and Kruskal-Wallis test (non-normal distribution) were applied to the data. If a main effect or interaction were found in the tests above, Šídák’s test, Dunnett’s T3 test or Dunn’s *post-hoc* multiple comparison corrections were applied. Repeated measures ANOVA or a Friedman test was used to compare multiple groups and parameters with repeated variables. When main effects were observed in above tests, Šídák’s or Dunn’s test were used for multiple comparisons corrections, respectively. Additionally, for comparison of two factors and the interaction between them, from multiple groups and parameters where one of the factors has repeated measurements, was performed using two-way ANOVA (no missing variables) or mixed-models ANOVA (Restricted maximum likelihood model, REML). The ANOVA tests were followed by Šídák’s multiple comparison test if main effects or interactions were found. The association between two groups or parameters was compared with either Pearson’s or Spearman’s tests. A binomial distribution test was performed to compare the probability of brain regions representing above-chance levels for specific parameters.

### Decision tree classifier model

#### Data normalization – subtracting the mean value for each brain regions pair per mouse

The data from two mice (total 14) were ignored as they had less than 40 recorded brain regions pairs (out of 99). The mean value of each pair was computed and subtracted for each mouse separately. This helped to reduce the variability of the measurements across mice and improved classification accuracy. To reduce over-representation of a single stimulus in the computation of the mean value for a pair, we first averaged the mean value per stimulus (for a specific mouse) and then subtracted the average of these means.

#### Averaging bouts

The average bout for each stimulus was computed for each session.

#### Data Imputation

For each mouse, a slightly different set of brain areas were recorded due to slight inaccuracies in placing the electrodes and slight difference in the individual mouse anatomy. This resulted in missing entries from some of the brain regions pairs. We used a data imputation strategy to restore these missing entries. Note that before this step, we subtracted the mean value per brain region per mouse and averaged all the bouts from the same stimuli of the same session. The imputation algorithm is based on the MICE algorithm [44]. It is an iterative algorithm. In each iteration, it estimates the missing entries by a linear combination of (some of) the other entries. The used data imputation algorithm was defined as follows:

1. For each missing value of brain pair i in bout b, (𝑏𝑝_𝑖,𝑏_), replace 𝑏𝑝_𝑖,𝑏_ with the average value of 𝑏𝑝_𝑖_ across the valid values 𝑏𝑝_𝑖_ (from all bouts of all mice with a valid measurement of 𝑏𝑝_𝑖_).
2. For each 𝑏𝑝_𝑖_ (𝑜𝑟𝑑𝑒𝑟 𝑜𝑓 𝑏𝑟𝑎𝑖𝑛 𝑝𝑎𝑖𝑟𝑠 𝑖𝑠 𝑟𝑎𝑛𝑑𝑜𝑚𝑖𝑧𝑒𝑑):

a. Randomly choose a set of predicting brain pairs {𝑏𝑝_𝑗_} such that 𝑏𝑝_𝑖_ ∉{𝑏𝑝_𝑗_} and |{𝑏𝑝_𝑗_}| < 0.5*number_of_equations. Where the number_of_equations equals to the number of (averaged) bouts from all the mice (66 predicting brain pairs as the number of bouts in our dataset is 131 average bouts).
b. Compute linear regression coefficients {aj} (by least square method) to minimize:

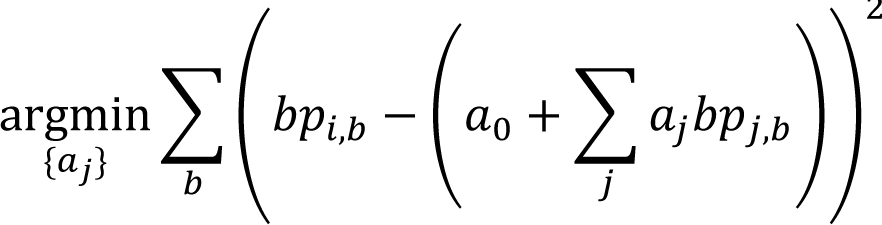
c. For each bout b in which 𝑏𝑝_𝑖,𝑏_ was not measured in-vivo, replace it with its estimation: 𝑎_0_ + ∑_𝑗_ 𝑎_𝑗_𝑏𝑝_𝑗,𝑏_
3. Repeat steps 2 for 20 iterations. Code was implemented in Matlab 2021a.

#### Classification and computation of confusion matrixes

We used Matlab’s *TreeBagger()* function to train a multi-class Random forest classifier for discriminating between a pair of stimuli (social vs object) or contexts (3 classes) or between stimuli (6 classes) or between. We used 80 random trees (a parameter of the TreeBagger function). We used cross-validation with “one mouse leave out” strategy to compute a confusion matrix for each mouse based on a training set that includes examples from all of the other mice. We balanced the training set to have the same number of examples from each class by randomly removing some of the training examples. Since both the balancing and the Random forest algorithm have a random component, we repeated the estimation of confusion matrixes 100 times (for each mouse) to better estimate the confusion matrixes. We then summed up all of the confusion matrixes (totally 1200 confusion matrixes: 12 mice and 100 confusion matrixes per mouse) and computed for each pair of classes (i,j) the percent of cases where the prediction was i for bouts of class j.

### Statistical Analysis

All tests were corrected for multiple comparisons using FDR corrections [45]. To compute p-values, we used the average (over 100 iterations) confusion matrix for each mouse (totally 12 confusion matrixes in which each cell i,j represents the % of predictions of class i for bouts of class j) and compare those with a set of random confusion matrixes which were generated by the same procedure except for replacing the trained classifier with a random classifier. This random classifier generated random labels with uniform distribution. To better describe the distribution of the random confusion matrixes, we generated 83 random confusion matrixes per mouse (each one of them is an average of 100 confusion matrixes). Then, the p-value for each cell in the confusion matrix was computed separately by comparing the 12 values from the confusion matrix of the trained classifier to the 83*12=996 values from the confusion matrixes of the random classifier using Mann-Whitney U test. In case a mouse did not have a bout from a specific class, this mouse was ignored in the computation of the p-value for the cells of this ground truth class.

## Author contributions

A.N.M.: Formal analysis, Investigation, Methodology, Validation, Visualization, Writing - original draft, and Writing - review & editing; D.P.: Formal analysis, and software.; S.N.: Data curation, Project administration, Software, Validation, Visualization, Writing - original draft, and Writing - review & editing. S.W.: Conceptualization, Funding acquisition, Project administration, Resources, Supervision, Writing - original draft, and Writing - review & editing

## Supporting information

See also Fig. S9.

## Acknowledgments

We thank Boris Shklyar, Head of the Bio-imaging Unit and Dr. Maya Lalzar, Head of the Bioinformatics Unit of the Faculty of Natural Sciences of the University of Haifa for their guidance and technical assistance. We also thank Eng. Alex Bizer, the experimental systems engineer of the Faculty of Natural Sciences of the University of Haifa, for their help.

## Brain-region name abbreviations

AcbC: Accumbens nucleus, core
AcbSh: Accumbens nucleus, shell
AhiAL: Amygdalo-hippocampal area, anterolateral part
BLA: Basolateral amygdaloid nucleus
BMP: Basomedial amygdaloid nucleus, posterior part
DMD: Dorsomedial hypothalamic nucleus, dorsal part
EA: Extended amygdala
IL: Infralimbic prefrontal cortex
LS: Lateral septum
MeAD: Medial amygdaloid nucleus, anterodorsal
Pir: Piriform cortex
PLH: Peduncular part of the lateral hypothalamus
PrL: Prelimbic prefrontal cortex
PVN: Paraventricular hypothalamic nucleus
vCA1: Field CA1 of the hippocampus, ventral part
vDG: Dentate gyrus, ventral part
VP: Ventral pallidum

## References

1. Adolphs R. Conceptual challenges and directions for social neuroscience. Neuron. 2010;65(6):752–67. Epub 2010/03/30. doi: 10.1016/j.neuron.2010.03.006. PubMed PMID: 20346753; PubMed Central PMCID: PMCPMC2887730.

2. Ford CL, Young LJ. Translational opportunities for circuit-based social neuroscience: advancing 21st century psychiatry. Curr Opin Neurobiol. 2021;68:1–8. Epub 2020/12/02. doi: 10.1016/j.conb.2020.11.007. PubMed PMID: 33260106; PubMed Central PMCID: PMCPMC8160019.

3. Kohl J, Dulac C. Neural control of parental behaviors. Curr Opin Neurobiol. 2018;49:116–22. Epub 2018/02/27. doi: 10.1016/j.conb.2018.02.002. PubMed PMID: 29482085; PubMed Central PMCID: PMCPMC6029232.

4. McKinsey G, Ahmed OM, Shah NM. Neural control of sexually dimorphic social behaviors. Curr Opin Physiol. 2018;6:89–95. Epub 2019/09/20. doi: 10.1016/j.cophys.2018.08.003. PubMed PMID: 31535059; PubMed Central PMCID: PMCPMC6750220.

5. Wei D, Talwar V, Lin D. Neural circuits of social behaviors: Innate yet flexible. Neuron. 2021;109(10):1600–20. Epub 2021/03/12. doi: 10.1016/j.neuron.2021.02.012. PubMed PMID: 33705708; PubMed Central PMCID: PMCPMC8141016.

6. Insel TR, Fernald RD. How the brain processes social information: searching for the social brain. Annu Rev Neurosci. 2004;27:697–722. Epub 2004/06/26. doi: 10.1146/annurev.neuro.27.070203.144148. PubMed PMID: 15217348.

7. Dickinson SY, Kelly DA, Padilla SL, Bergan JF. From Reductionism Toward Integration: Understanding How Social Behavior Emerges From Integrated Circuits. Front Integr Neurosci. 2022;16:862437. Epub 2022/04/19. doi: 10.3389/fnint.2022.862437. PubMed PMID: 35431824; PubMed Central PMCID: PMCPMC9010670.

8. Goodson JL. The vertebrate social behavior network: evolutionary themes and variations. Horm Behav. 2005;48(1):11–22. Epub 2005/05/12. doi: 10.1016/j.yhbeh.2005.02.003. PubMed PMID: 15885690; PubMed Central PMCID: PMCPMC2570781.

9. Dai B, Sun F, Tong X, Ding Y, Kuang A, Osakada T, et al. Responses and functions of dopamine in nucleus accumbens core during social behaviors. Cell Rep. 2022;40(8):111246. Epub 2022/08/25. doi: 10.1016/j.celrep.2022.111246. PubMed PMID: 36001967; PubMed Central PMCID: PMCPMC9511885.

10. Felix-Ortiz AC, Tye KM. Amygdala inputs to the ventral hippocampus bidirectionally modulate social behavior. J Neurosci. 2014;34(2):586–95. Epub 2014/01/10. doi: 10.1523/JNEUROSCI.4257-13.2014. PubMed PMID: 24403157; PubMed Central PMCID: PMCPMC3870937.

11. Kingsbury L, Huang S, Raam T, Ye LS, Wei D, Hu RK, et al. Cortical Representations of Conspecific Sex Shape Social Behavior. Neuron. 2020;107(5):941–53 e7. Epub 2020/07/15. doi: 10.1016/j.neuron.2020.06.020. PubMed PMID: 32663438; PubMed Central PMCID: PMCPMC7486272.

12. Leroy F, Park J, Asok A, Brann DH, Meira T, Boyle LM, et al. A circuit from hippocampal CA2 to lateral septum disinhibits social aggression. Nature. 2018;564(7735):213–8. Epub 2018/12/07. doi: 10.1038/s41586-018-0772-0. PubMed PMID: 30518859; PubMed Central PMCID: PMCPMC6364572.

13. Kohl J, Babayan BM, Rubinstein ND, Autry AE, Marin-Rodriguez B, Kapoor V, et al. Functional circuit architecture underlying parental behaviour. Nature. 2018;556(7701):326–31. Epub 2018/04/13. doi: 10.1038/s41586-018-0027-0. PubMed PMID: 29643503; PubMed Central PMCID: PMCPMC5908752.

14. Scheggia D, La Greca F, Maltese F, Chiacchierini G, Italia M, Molent C, et al. Reciprocal cortico-amygdala connections regulate prosocial and selfish choices in mice. Nat Neurosci. 2022;25(11):1505–18. Epub 2022/10/26. doi: 10.1038/s41593-022-01179-2. PubMed PMID: 36280797; PubMed Central PMCID: PMCPMC7613781.

15. Swanson LW, Hahn JD, Sporns O. Structure-function subsystem models of female and male forebrain networks integrating cognition, affect, behavior, and bodily functions. Proc Natl Acad Sci U S A. 2020;117(49):31470–81. Epub 2020/11/25. doi: 10.1073/pnas.2017733117. PubMed PMID: 33229546; PubMed Central PMCID: PMCPMC7733829.

16. Twining RC, Vantrease JE, Love S, Padival M, Rosenkranz JA. An intra-amygdala circuit specifically regulates social fear learning. Nat Neurosci. 2017;20(3):459–69. Epub 2017/01/24. doi: 10.1038/nn.4481. PubMed PMID: 28114293; PubMed Central PMCID: PMCPMC5323274.

17. Yamamoto R, Ahmed N, Ito T, Gungor NZ, Pare D. Optogenetic Study of Anterior BNST and Basomedial Amygdala Projections to the Ventromedial Hypothalamus. Eneuro. 2018;5(3). Epub 2018/07/05. doi: 10.1523/ENEURO.0204-18.2018. PubMed PMID: 29971248; PubMed Central PMCID: PMCPMC6027956.

18. Dolen G, Darvishzadeh A, Huang KW, Malenka RC. Social reward requires coordinated activity of nucleus accumbens oxytocin and serotonin. Nature. 2013;501(7466):179–84. doi: 10.1038/nature12518. PubMed PMID: 24025838; PubMed Central PMCID: PMCPMC4091761.

19. Felix-Ortiz AC, Burgos-Robles A, Bhagat ND, Leppla CA, Tye KM. Bidirectional modulation of anxiety-related and social behaviors by amygdala projections to the medial prefrontal cortex. Neuroscience. 2016;321:197–209. Epub 2015/07/25. doi: 10.1016/j.neuroscience.2015.07.041. PubMed PMID: 26204817; PubMed Central PMCID: PMCPMC4721937.

20. Huang WC, Zucca A, Levy J, Page DT. Social Behavior Is Modulated by Valence-Encoding mPFC-Amygdala Sub-circuitry. Cell Rep. 2020;32(2):107899. Epub 2020/07/16. doi: 10.1016/j.celrep.2020.107899. PubMed PMID: 32668253; PubMed Central PMCID: PMCPMC7410267.

21. Reis FM, Lee JY, Maesta-Pereira S, Schuette PJ, Chakerian M, Liu J, et al. Dorsal periaqueductal gray ensembles represent approach and avoidance states. Elife. 2021;10. Epub 2021/05/07. doi: 10.7554/eLife.64934. PubMed PMID: 33955356; PubMed Central PMCID: PMCPMC8133778.

22. Mague SD, Talbot A, Blount C, Walder-Christensen KK, Duffney LJ, Adamson E, et al. Brain-wide electrical dynamics encode individual appetitive social behavior. Neuron. 2022;110(10):1728–41 e7. Epub 2022/03/17. doi: 10.1016/j.neuron.2022.02.016. PubMed PMID: 35294900; PubMed Central PMCID: PMCPMC9126093.

23. Schaich Borg J, Srivastava S, Lin L, Heffner J, Dunson D, Dzirasa K, et al. Rat intersubjective decisions are encoded by frequency-specific oscillatory contexts. Brain Behav. 2017;7(6):e00710. Epub 2017/06/24. doi: 10.1002/brb3.710. PubMed PMID: 28638715; PubMed Central PMCID: PMCPMC5474713.

24. Buzsaki G, Draguhn A. Neuronal oscillations in cortical networks. Science. 2004;304(5679):1926–9. doi: 10.1126/science.1099745. PubMed PMID: 15218136.

25. Harris AZ, Gordon JA. Long-range neural synchrony in behavior. Annu Rev Neurosci. 2015;38:171–94. Epub 2015/04/22. doi: 10.1146/annurev-neuro-071714-034111. PubMed PMID: 25897876; PubMed Central PMCID: PMCPMC4497851.

26. Uhlhaas PJ, Pipa G, Lima B, Melloni L, Neuenschwander S, Nikolic D, et al. Neural synchrony in cortical networks: history, concept and current status. Front Integr Neurosci. 2009;3:17. Epub 2009/08/12. doi: 10.3389/neuro.07.017.2009. PubMed PMID: 19668703; PubMed Central PMCID: PMCPMC2723047.

27. Bocchio M, Nabavi S, Capogna M. Synaptic Plasticity, Engrams, and Network Oscillations in Amygdala Circuits for Storage and Retrieval of Emotional Memories. Neuron. 2017;94(4):731–43. Epub 2017/05/19. doi: 10.1016/j.neuron.2017.03.022. PubMed PMID: 28521127.

28. Chen S, Tan Z, Xia W, Gomes CA, Zhang X, Zhou W, et al. Theta oscillations synchronize human medial prefrontal cortex and amygdala during fear learning. Sci Adv. 2021;7(34). Epub 2021/08/20. doi: 10.1126/sciadv.abf4198. PubMed PMID: 34407939; PubMed Central PMCID: PMCPMC8373137.

29. Taub AH, Perets R, Kahana E, Paz R. Oscillations Synchronize Amygdala-to-Prefrontal Primate Circuits during Aversive Learning. Neuron. 2018;97(2):291–8 e3. Epub 2018/01/02. doi: 10.1016/j.neuron.2017.11.042. PubMed PMID: 29290553.

30. John SR, Dagash W, Mohapatra AN, Netser S, Wagner S. Distinct Dynamics of Theta and Gamma Rhythmicity during Social Interaction Suggest Differential Mode of Action in the Medial Amygdala of Sprague Dawley Rats and C57BL/6J Mice. Neuroscience. 2022;493:69–80. Epub 2022/05/02. doi: 10.1016/j.neuroscience.2022.04.020. PubMed PMID: 35490969.

31. Kuga N, Abe R, Takano K, Ikegaya Y, Sasaki T. Prefrontal-amygdalar oscillations related to social behavior in mice. Elife. 2022;11. Epub 2022/05/18. doi: 10.7554/eLife.78428. PubMed PMID: 35580019; PubMed Central PMCID: PMCPMC9113747.

32. Tendler A, Wagner S. Different types of theta rhythmicity are induced by social and fearful stimuli in a network associated with social memory. Elife. 2015;4. doi: 10.7554/eLife.03614. PubMed PMID: 25686218; PubMed Central PMCID: PMCPMC4353977.

33. Buzsaki G, Watson BO. Brain rhythms and neural syntax: implications for efficient coding of cognitive content and neuropsychiatric disease. Dialogues Clin Neurosci. 2012;14(4):345–67. Epub 2013/02/09. doi: 10.31887/DCNS.2012.14.4/gbuzsaki. PubMed PMID: 23393413; PubMed Central PMCID: PMCPMC3553572.

34. Cannon J, McCarthy MM, Lee S, Lee J, Borgers C, Whittington MA, et al. Neurosystems: brain rhythms and cognitive processing. Eur J Neurosci. 2014;39(5):705–19. Epub 2013/12/18. doi: 10.1111/ejn.12453. PubMed PMID: 24329933; PubMed Central PMCID: PMCPMC4916881.

35. Uhlhaas PJ, Roux F, Rodriguez E, Rotarska-Jagiela A, Singer W. Neural synchrony and the development of cortical networks. Trends Cogn Sci. 2010;14(2):72–80. doi: 10.1016/j.tics.2009.12.002. PubMed PMID: 20080054.

36. Geschwind DH, Levitt P. Autism spectrum disorders: developmental disconnection syndromes. Curr Opin Neurobiol. 2007;17(1):103–11. Epub 2007/02/06. doi: 10.1016/j.conb.2007.01.009. PubMed PMID: 17275283.

37. Lazaro MT, Taxidis J, Shuman T, Bachmutsky I, Ikrar T, Santos R, et al. Reduced Prefrontal Synaptic Connectivity and Disturbed Oscillatory Population Dynamics in the CNTNAP2 Model of Autism. Cell Rep. 2019;27(9):2567–78 e6. Epub 2019/05/30. doi: 10.1016/j.celrep.2019.05.006. PubMed PMID: 31141683; PubMed Central PMCID: PMCPMC6553483.

38. Fries P. Rhythms for Cognition: Communication through Coherence. Neuron. 2015;88(1):220–35. Epub 2015/10/09. doi: 10.1016/j.neuron.2015.09.034. PubMed PMID: 26447583; PubMed Central PMCID: PMCPMC4605134.

39. Chen S, He L, Huang AJY, Boehringer R, Robert V, Wintzer ME, et al. A hypothalamic novelty signal modulates hippocampal memory. Nature. 2020;586(7828):270–4. Epub 2020/10/02. doi: 10.1038/s41586-020-2771-1. PubMed PMID: 32999460.

40. McHugh TJ, Jones MW, Quinn JJ, Balthasar N, Coppari R, Elmquist JK, et al. Dentate gyrus NMDA receptors mediate rapid pattern separation in the hippocampal network. Science. 2007;317(5834):94–9. Epub 2007/06/09. doi: 10.1126/science.1140263. PubMed PMID: 17556551.

41. Mohapatra AN, Netser S, Wagner S. Modular Electrode Array for Multi-site Extracellular Recordings from Brains of Freely Moving Rodents. Curr Protoc. 2022;2(5):e399. Epub 2022/05/11. doi: 10.1002/cpz1.399. PubMed PMID: 35536117.

42. Netser S, Haskal S, Magalnik H, Bizer A, Wagner S. A System for Tracking the Dynamics of Social Preference Behavior in Small Rodents. J Vis Exp. 2019;(153). doi: 10.3791/60336. PubMed PMID: 31814614.

43. Barnett L, Seth AK. The MVGC multivariate Granger causality toolbox: a new approach to Granger-causal inference. J Neurosci Methods. 2014;223:50–68. Epub 2013/11/10. doi: 10.1016/j.jneumeth.2013.10.018. PubMed PMID: 24200508.

44. Azur MJ, Stuart EA, Frangakis C, Leaf PJ. Multiple imputation by chained equations: what is it and how does it work? Int J Methods Psychiatr Res. 2011;20(1):40–9. Epub 2011/04/19. doi: 10.1002/mpr.329. PubMed PMID: 21499542; PubMed Central PMCID: PMCPMC3074241.

45. Boca SM, Leek JT. A direct approach to estimating false discovery rates conditional on covariates. PeerJ. 2018;6:e6035. Epub 2018/12/26. doi: 10.7717/peerj.6035. PubMed PMID: 30581661; PubMed Central PMCID: PMCPMC6292380.

46. Jabarin R, Levy N, Abergel Y, Berman JH, Zag A, Netser S, et al. Pharmacological modulation of AMPA receptors rescues specific impairments in social behavior associated with the A350V Iqsec2 mutation. Transl Psychiatry. 2021;11(1):234. Epub 2021/04/24. doi: 10.1038/s41398-021-01347-1. PubMed PMID: 33888678; PubMed Central PMCID: PMCPMC8062516.

47. Netser S, Meyer A, Magalnik H, Zylbertal A, de la Zerda SH, Briller M, et al. Distinct dynamics of social motivation drive differential social behavior in laboratory rat and mouse strains. Nat Commun. 2020;11(1):5908. Epub 2020/11/22. doi: 10.1038/s41467-020-19569-0. PubMed PMID: 33219219; PubMed Central PMCID: PMCPMC7679456.

48. Clayton MS, Yeung N, Kadosh RC. The roles of cortical oscillations in sustained attention. Trends in Cognitive Sciences. 2015;19(4):188–95. doi: 10.1016/j.tics.2015.02.004. PubMed PMID: WOS:000352674600006.

49. Fiebelkorn IC, Kastner S. A Rhythmic Theory of Attention. Trends in Cognitive Sciences. 2019;23(2):87–101. doi: 10.1016/j.tics.2018.11.009. PubMed PMID: WOS:000455717200005.

50. Helfrich RF, Breska A, Knight RT. Neural entrainment and network resonance in support of top-down guided attention. Curr Opin Psychol. 2019;29:82–9. Epub 2019/01/29. doi: 10.1016/j.copsyc.2018.12.016. PubMed PMID: 30690228; PubMed Central PMCID: PMCPMC6606401.

51. Karakas S. A review of theta oscillation and its functional correlates. Int J Psychophysiol. 2020;157:82–99. Epub 2020/05/20. doi: 10.1016/j.ijpsycho.2020.04.008. PubMed PMID: 32428524.

52. Knyazev GG. Motivation, emotion, and their inhibitory control mirrored in brain oscillations. Neurosci Biobehav R. 2007;31(3):377–95. doi: 10.1016/j.neubiorev.2006.10.004. PubMed PMID: WOS:000246316100006.

53. Buzsaki G, Wang XJ. Mechanisms of gamma oscillations. Annu Rev Neurosci. 2012;35:203–25. Epub 2012/03/27. doi: 10.1146/annurev-neuro-062111-150444. PubMed PMID: 22443509; PubMed Central PMCID: PMCPMC4049541.

54. Headley DB, Pare D. In sync: gamma oscillations and emotional memory. Front Behav Neurosci. 2013;7:170. Epub 2013/12/10. doi: 10.3389/fnbeh.2013.00170. PubMed PMID: 24319416; PubMed Central PMCID: PMCPMC3836200.

55. Benchenane K, Tiesinga PH, Battaglia FP. Oscillations in the prefrontal cortex: a gateway to memory and attention. Curr Opin Neurobiol. 2011;21(3):475–85. Epub 2011/03/25. doi: 10.1016/j.conb.2011.01.004. PubMed PMID: 21429736.

56. Palva JM, Palva S. Functional integration across oscillation frequencies by cross-frequency phase synchronization. Eur J Neurosci. 2018;48(7):2399–406. Epub 2017/11/03. doi: 10.1111/ejn.13767. PubMed PMID: 29094462.

57. Cai Y, Tang X, Chen X, Li X, Wang Y, Bao X, et al. Liver X receptor beta regulates the development of the dentate gyrus and autistic-like behavior in the mouse. Proc Natl Acad Sci U S A. 2018;115(12):E2725–E33. Epub 2018/03/07. doi: 10.1073/pnas.1800184115. PubMed PMID: 29507213; PubMed Central PMCID: PMCPMC5866608.

58. Doucette E, Merfeld E, Leblanc H, Monasterio A, Cincotta C, Grella SL, et al. Social behavior in mice following chronic optogenetic stimulation of hippocampal engrams. Neurobiol Learn Mem. 2020;176:107321. Epub 2020/11/10. doi: 10.1016/j.nlm.2020.107321. PubMed PMID: 33164892; PubMed Central PMCID: PMCPMC7708439.

59. Leung C, Cao F, Nguyen R, Joshi K, Aqrabawi AJ, Xia S, et al. Activation of Entorhinal Cortical Projections to the Dentate Gyrus Underlies Social Memory Retrieval. Cell Rep. 2018;23(8):2379–91. Epub 2018/05/24. doi: 10.1016/j.celrep.2018.04.073. PubMed PMID: 29791849.

60. Cope EC, Waters RC, Diethorn EJ, Pagliai KA, Dias CG, Tsuda M, et al. Adult-Born Neurons in the Hippocampus Are Essential for Social Memory Maintenance. Eneuro. 2020;7(6). Epub 2020/10/17. doi: 10.1523/ENEURO.0182-20.2020. PubMed PMID: 33060182; PubMed Central PMCID: PMCPMC7768285.

61. Li J, Sun X, You Y, Li Q, Wei C, Zhao L, et al. Auts2 deletion involves in DG hypoplasia and social recognition deficit: The developmental and neural circuit mechanisms. Sci Adv. 2022;8(9):eabk1238. Epub 2022/03/03. doi: 10.1126/sciadv.abk1238. PubMed PMID: 35235353; PubMed Central PMCID: PMCPMC8890717.

62. Leutgeb JK, Leutgeb S, Moser MB, Moser EI. Pattern separation in the dentate gyrus and CA3 of the hippocampus. Science. 2007;315(5814):961–6. Epub 2007/02/17. doi: 10.1126/science.1135801. PubMed PMID: 17303747.

63. Wu WY, Yiu E, Ophir AG, Smith DM. Effects of social context manipulation on dorsal and ventral hippocampal neuronal responses. Hippocampus. 2023. Epub 2023/02/16. doi: 10.1002/hipo.23507. PubMed PMID: 36789678.

