## Supplementary material for "Synchronized LFP rhythmicity in the social brain reflects the context of social encounters": See also Fig. S9.

### Supplementary figure legends

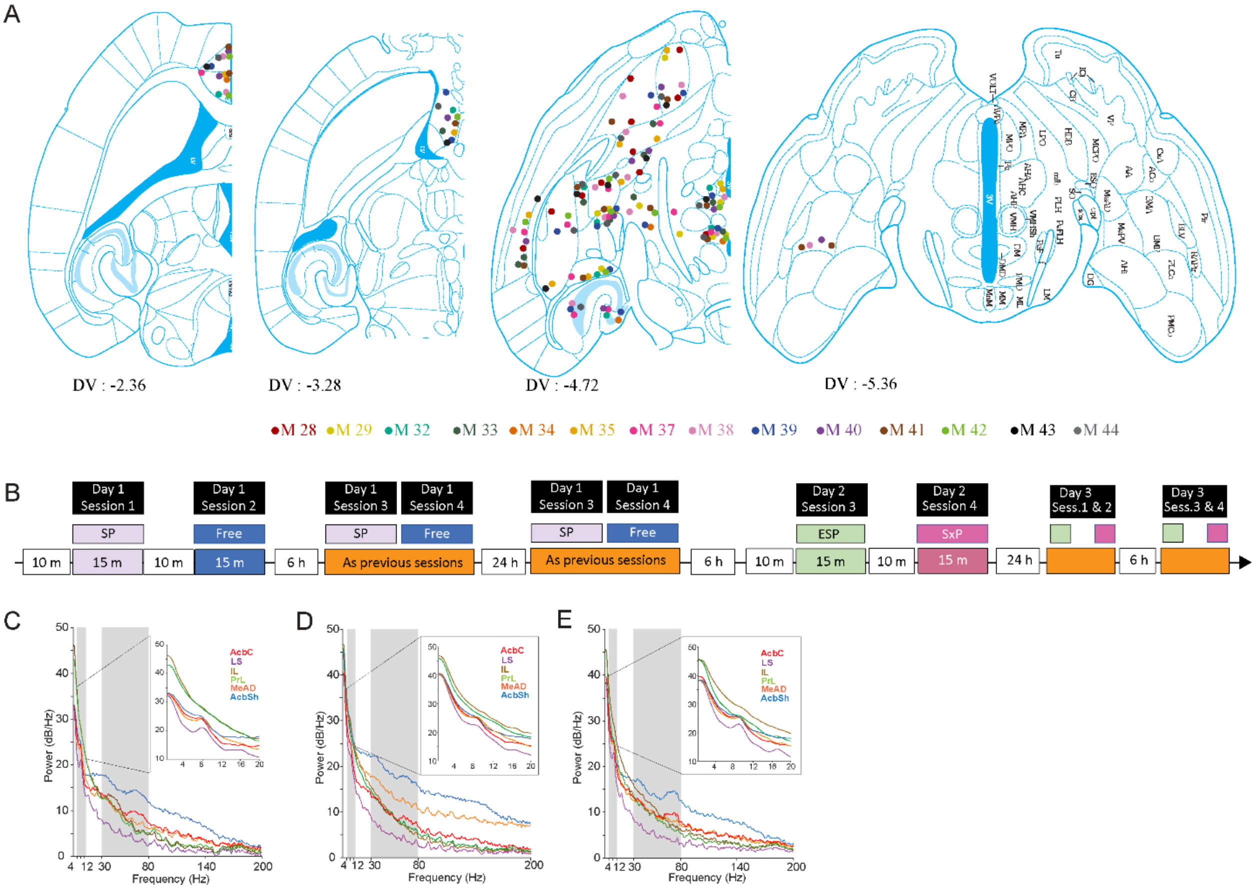


**Figure S1: Locations of all electrode tips and PSD profiles from individual sessions**

A. The locations of all electrode tips, as verified post mortem, color-coded for all subject animals (M28-44) on brain atlas pictures. The legend of the color code is presented below.

B. Timeline of all sessions conducted by each subject.

C. The power spectral density (PSD) plots of LFP signals recorded in the MeAD, IL, PrL, CA1, AcbC, and AcbSh during the encounter period of SP (left), EsP (middle) and SxP (right) task single sessions, all conducted by the same mouse. The inset shows the PSD plots for 0-20 Hz band at higher resolution.

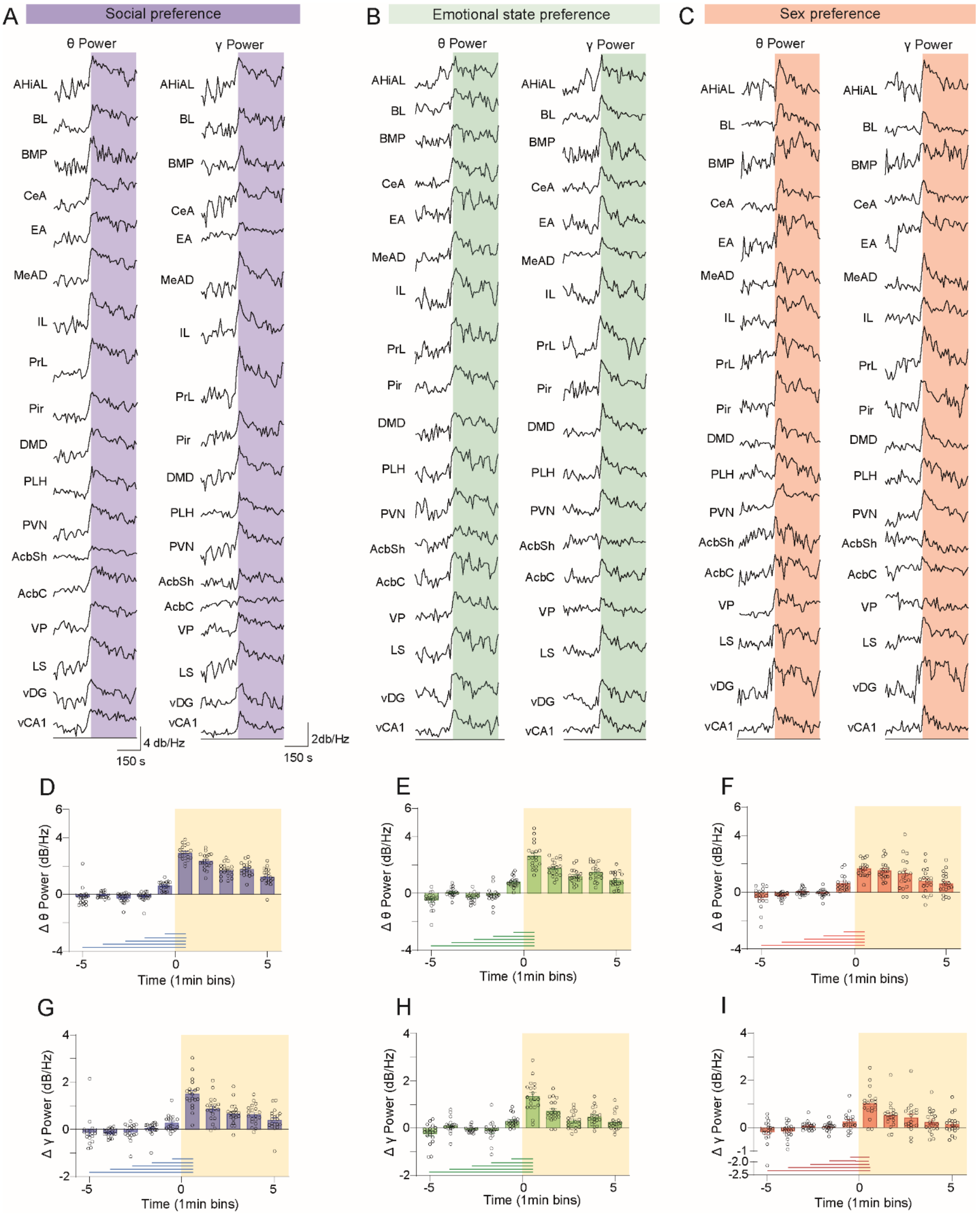
A
Social preference
θ Power
γ Power
AHiAL
BL
BMP
CeA
EA
MeAD
IL
PrL
Pir
DMD
PLH
PVN
AcbSh
AcbC
VP
LS
vDG
vCA1
4 db/Hz
150 s
2db/Hz
150 s
B
Emotional state preference
C
Sex preference
D
E
F
Δ θ Power (dB/Hz)
Time (1min bins)
G
H
I
Δ γ Power (dB/Hz)
Time (1min bins)

**Figure S2. Similar dynamics of theta and gamma power across the various tasks and brain regions**

A. Mean traces of ΔθP (left column) and ΔγP (right column) across all sessions of the SP task for each brain region. The colored bar represents the encounter period.
B. As in A, for the EsP task
C. As in A, for the SxP task
D. Mean (±SEM) ΔθP, averaged across all brain regions for every minute of the baseline and encounters periods of the SP task. Time 0 min represents the time of stimuli insertion. Lines below the bars represent a significant difference between the first minute of the encounter and every minute of the baseline period.
E. As in D, for the EsP task.
F. As in D, for the SxP task

G-I, as in D-F, for ΔγP.

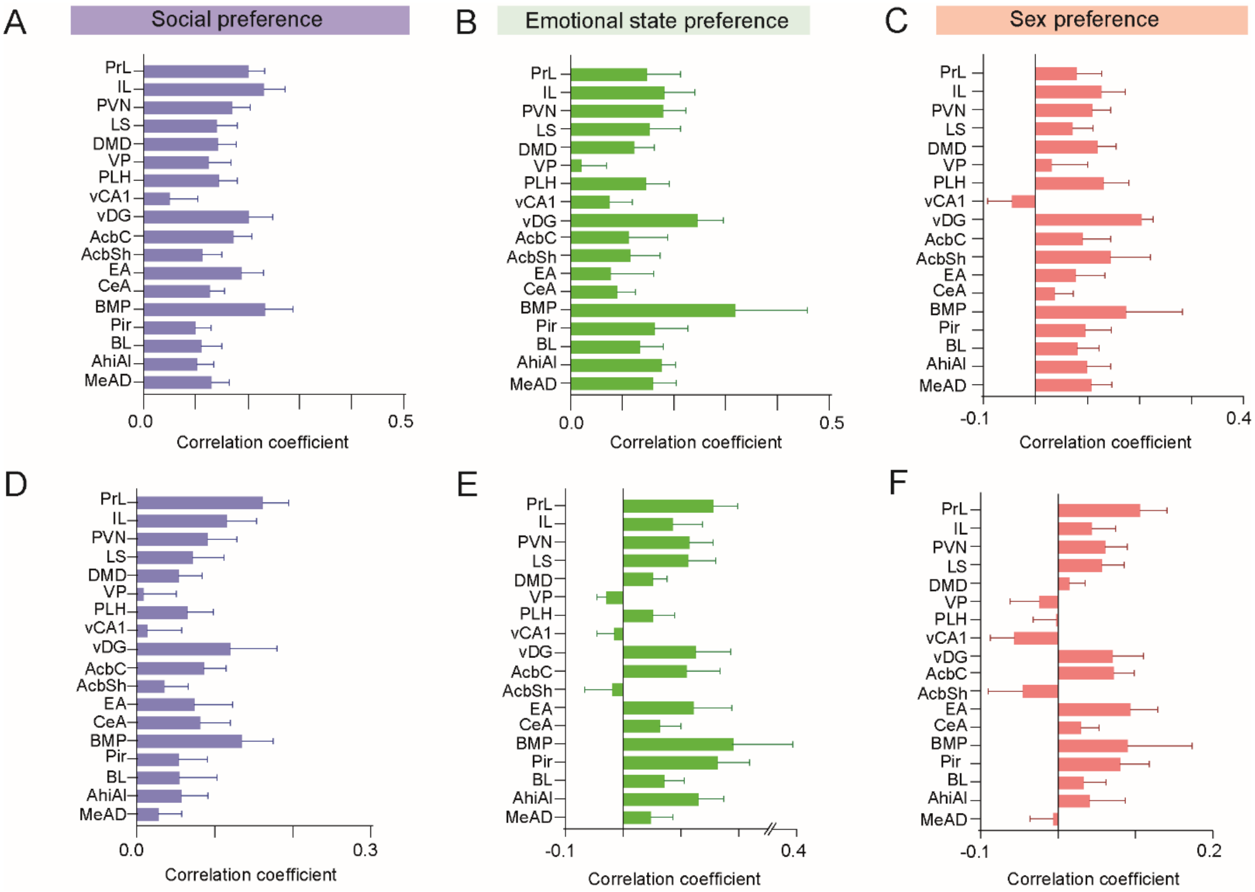


**Figure S3. No correlation exists between theta and gamma power changes during social encounters and subject speed**

A. Correlation coefficients across all SP task sessions for calculating Pearson's correlation between the change in theta power (ΔθP) and the mean speed of the subject.

B. As in A, for EsP sessions.

C. As in A, for SxP sessions.

D-F. As in A-C, for gamma power (ΔγP).

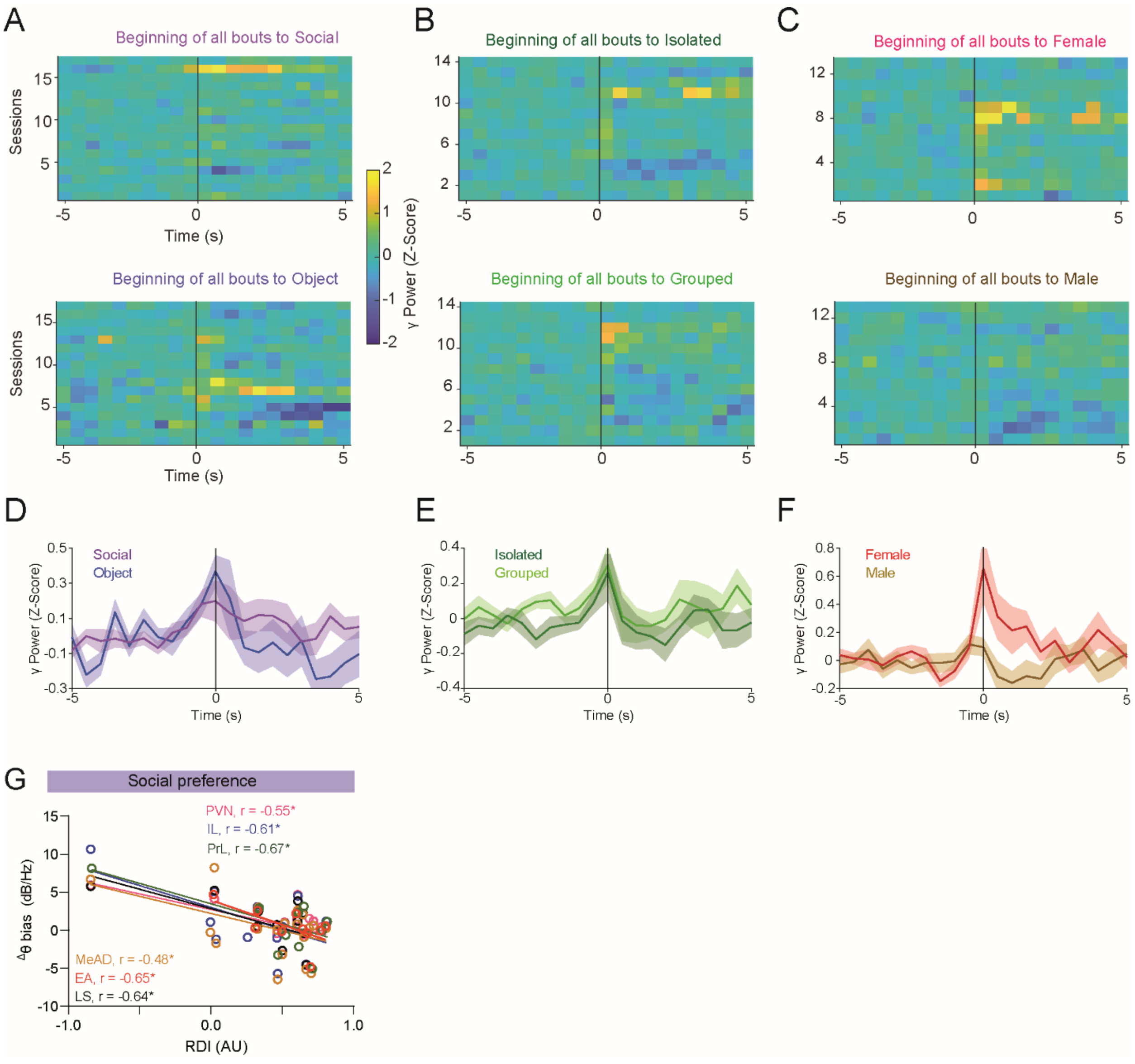


**Figure S4. Z-score analysis of changes in gamma power during social investigation of specific stimuli**

A. Heat maps of average gamma power in AhiAl, before and during social investigation bouts of subjects with social (above) and object (below) stimuli. Each row represents the mean Z-score of all bouts in a single session (using 0.5 s bins). Time '0' represents the beginning of the bout. The color code scale is on the right.

B. As in A, for investigation bouts of isolated (above) and grouped (below) social stimuli during EsP tasks.

C. As in A, for investigation bouts of female (above) and male (below) social stimuli during SxP tasks.

D. Mean (±SEM) Z-score trace of the data shown in A for both stimuli.

E. As in D. for the data shown in B.

F. As in D, for the data shown in C.

G. Correlation between mean change in theta power during investigation bouts ($^{\Delta}\theta$P) in specific brain regions and RDI values during the SP task, with no session excluded. Only statistically significant linear correlations are shown.

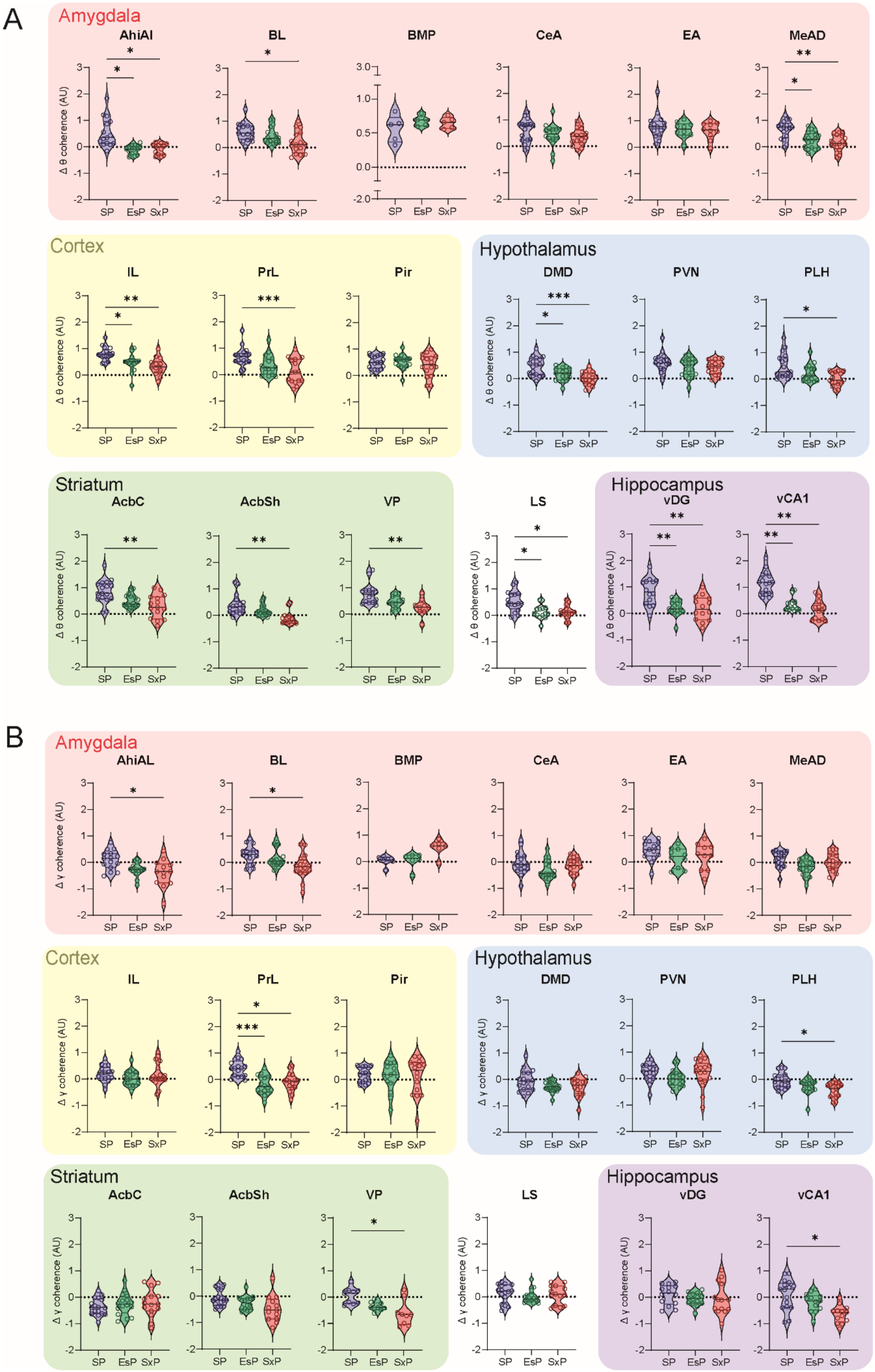
A
Amygdala
AhiAl
BL
BMP
CeA
EA
MeAD
Δ θ coherence (AU)
SP
EsP
SxP
Cortex
IL
PrL
Pir
Hypothalamus
DMD
PVN
PLH
Striatum
AcbC
AcbSh
VP
LS
Hippocampus
vDG
vCA1
B
Amygdala
AhiAL
BL
BMP
CeA
EA
MeAD
Δ γ coherence (AU)
Cortex
IL
PrL
Pir
Hypothalamus
DMD
PVN
PLH
Striatum
AcbC
AcbSh
VP
LS
Hippocampus
vDG
vCA1

**Figure S5. Coherence changes during the encounter period of the various tasks are brain region-specific**

A. Mean change in theta coherence (ΔθCo) between a given brain region and all other simultaneously recorded regions during the encounter period, compared across the three tasks.

B. As in A, for gamma coherence (ΔγCo).

*p<0.05, **p<0.01, ***p<0.001, ****p<0.0001, Dunnett's and Dunn's *post-hoc* test after FDR correction, following the main effect in ANOVA and Kruskal Wallis test, respectively.

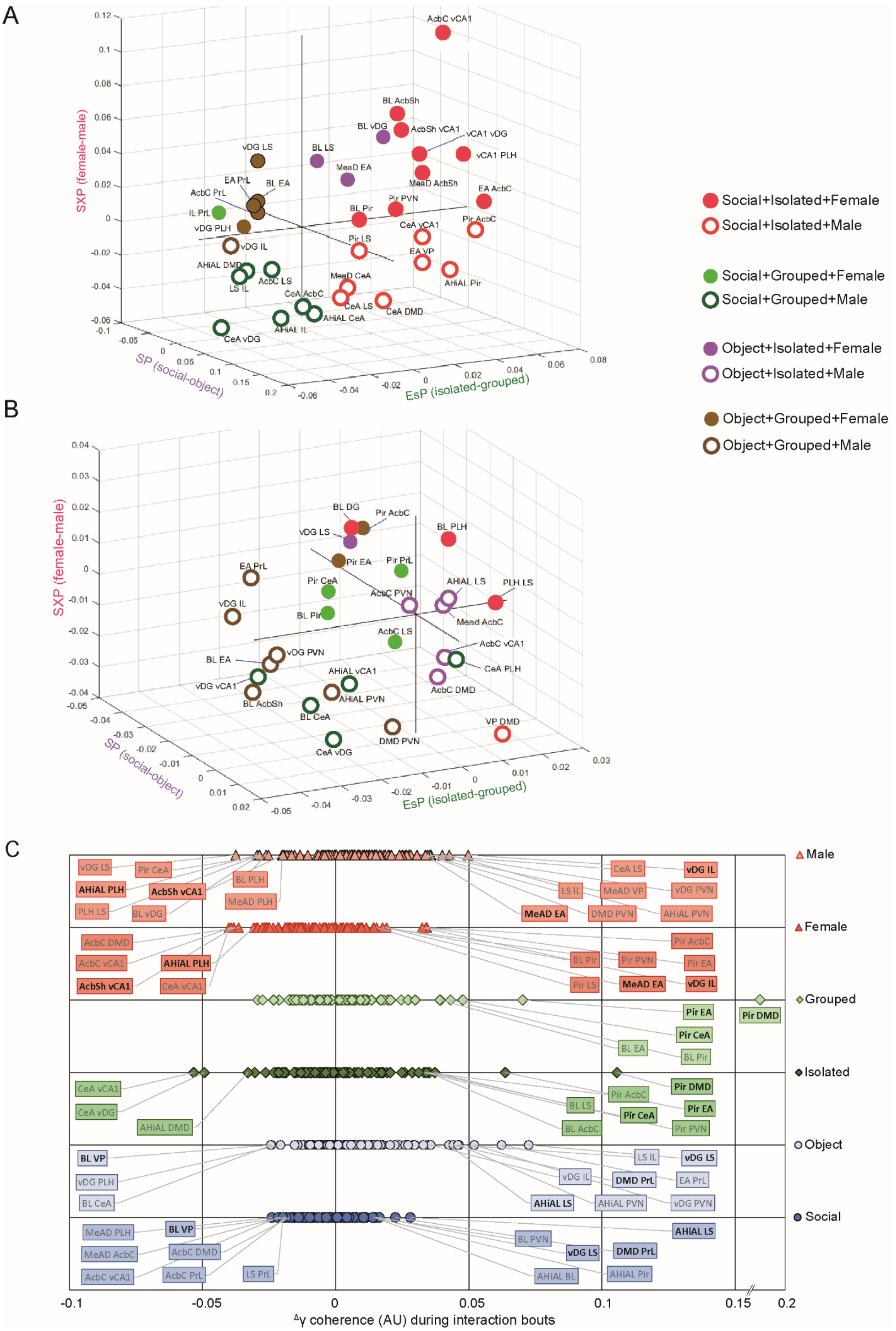
A
SXP (female-male)
SP (social-object)
EsP (isolated-grouped)
AcbC vCA1
BL AcbSh
BL vDG
AcbSh vCA1
vCA1 vDG
vCA1 PLH
vDG LS
BL LS
MeaD EA
MeaD AcbSh
EA AcbC
EA PrL
BL EA
AcbC PrL
IL PrL
vDG PLH
vDG IL
BL Pir
Pir PVN
CeA vCA1
Pir AcbC
Pir LS
EA VP
AHiAL DMD
LS IL
AcbC LS
MeaD CeA
AHiAL Pir
CeA AcbC
CeA LS
CeA DMD
AHiAL CeA
AHiAL IL
CeA vDG
B
BL DG
Pir AcbC
vDG LS
BL PLH
EA PrL
Pir EA
Pir PrL
Pir CeA
AHiAL LS
PLH LS
AcbC PVN
vDG IL
BL Pir
Mead AcbC
AcbC LS
AcbC vCA1
BL EA
vDG PVN
CeA PLH
AHiAL vCA1
vDG vCA1
AcbC DMD
AHiAL PVN
BL AcbSh
BL CeA
VP DMD
DMD PVN
CeA vDG
C
Male
Female
Grouped
Isolated
Object
Social
vDG LS
Pir CeA
AHiAL PLH
AcbSh vCA1
BL PLH
PLH LS
BL vDG
MeAD PLH
CeA LS
vDG IL
LS IL
MeAD VP
vDG PVN
MeAD EA
DMD PVN
AHiAL PVN
AcbC DMD
AcbC vCA1
AHiAL PLH
AcbSh vCA1
CeA vCA1
Pir AcbC
BL Pir
Pir PVN
Pir EA
Pir LS
MeAD EA
vDG IL
Pir EA
Pir DMD
Pir CeA
BL EA
BL Pir
CeA vCA1
CeA vDG
AHiAL DMD
Pir DMD
Pir AcbC
BL LS
Pir EA
Pir CeA
BL AcbC
Pir PVN
BL VP
vDG PLH
BL CeA
LS IL
vDG LS
vDG IL
DMD PrL
EA PrL
AHiAL LS
AHiAL PVN
vDG PVN
MeAD PLH
BL VP
MeAD AcbC
AcbC DMD
AcbC vCA1
AcbC PrL
LS PrL
BL PVN
AHiAL LS
vDG LS
DMD PrL
AHiAL BL
AHiAL Pir
-0.1
-0.05
0
0.05
0.1
0.15
0.2
Δγ coherence (AU) during interaction bouts
Social+Isolated+Female
Social+Isolated+Male
Social+Grouped+Female
Social+Grouped+Male
Object+Isolated+Female
Object+Isolated+Male
Object+Grouped+Female
Object+Grouped+Male

**Figure S6. Distribution of theta and gamma coherence changes during social investigation bouts across tasks**

A. A 3D representation of the mean difference in theta coherence change during investigation bouts ($^{\Delta}\theta$Co) between the preferred and less-preferred stimuli, across all tasks. Each dot represents a pair of brain regions (n = 84 pairs), color- and shape-coded according to the combined bias across all tasks. Only pairs which showed a coherence change of 1.5*SD from the mean are presented. We excluded any pair with fewer than five sessions of investigation bouts with one of the stimuli or with both.

B. As in A, for gamma coherence change ($^{\Delta}\gamma$Co)

C. Distribution of gamma coherence change ($\Delta\gamma$Co) between each pair of brain regions during investigation bouts, plotted separately for each stimulus used in the SP (blue), EsP (green) and SxP (red) tasks. The brain region pairs which passed the mean ± 1.5*SD cutoff are labeled and those that are shared by both stimuli in the same task are denoted in bold.

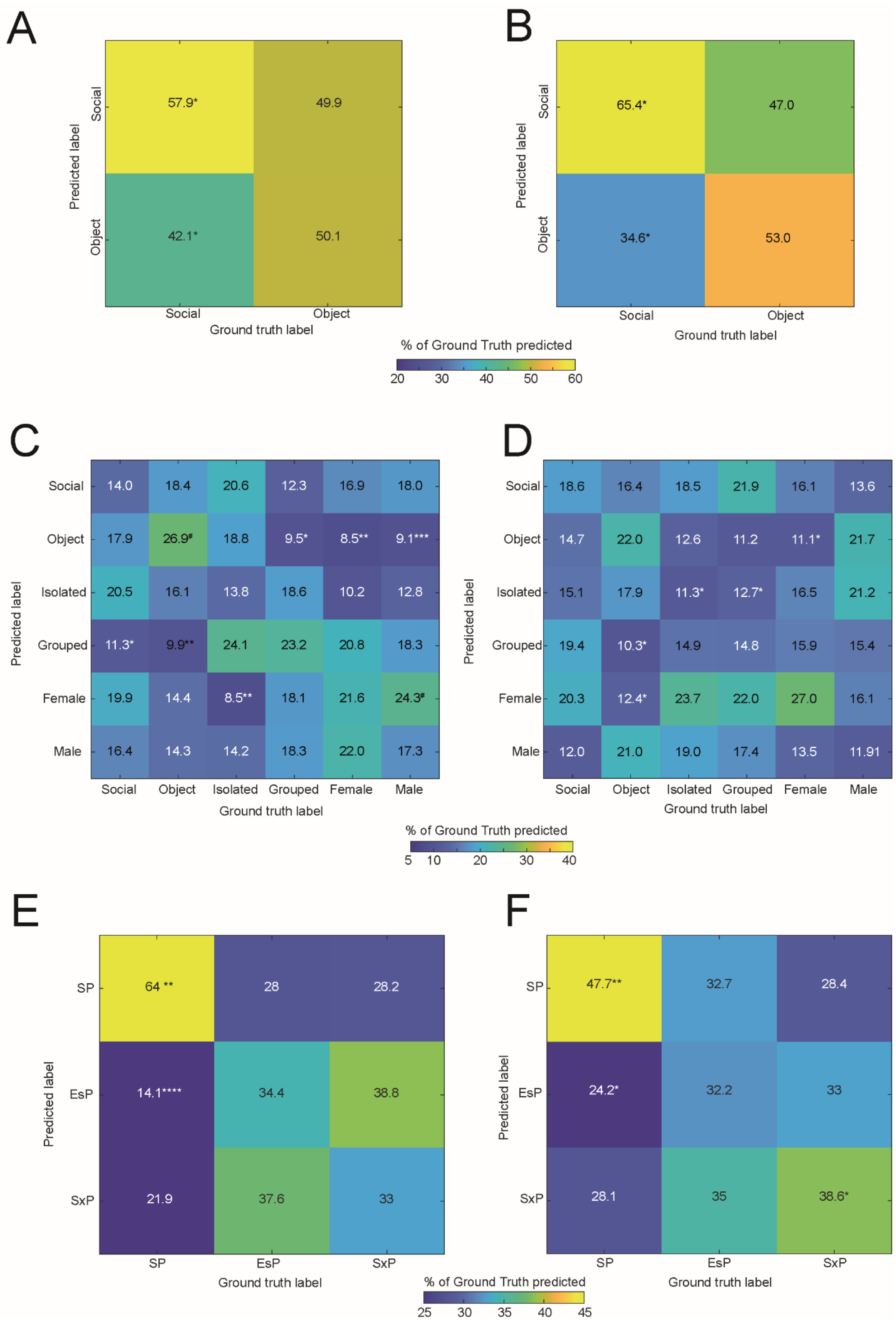
A
Social
Object
Predicted label
57.9*
49.9
42.1*
50.1
Social
Object
Ground truth label
B
65.4*
47.0
34.6*
53.0
% of Ground Truth predicted
20 30 40 50 60
C
Social Object Isolated Grouped Female Male
14.0 18.4 20.6 12.3 16.9 18.0
17.9 26.9# 18.8 9.5* 8.5** 9.1***
20.5 16.1 13.8 18.6 10.2 12.8
11.3* 9.9** 24.1 23.2 20.8 18.3
19.9 14.4 8.5** 18.1 21.6 24.3#
16.4 14.3 14.2 18.3 22.0 17.3
D
18.6 16.4 18.5 21.9 16.1 13.6
14.7 22.0 12.6 11.2 11.1* 21.7
15.1 17.9 11.3* 12.7* 16.5 21.2
19.4 10.3* 14.9 14.8 15.9 15.4
20.3 12.4* 23.7 22.0 27.0 16.1
12.0 21.0 19.0 17.4 13.5 11.91
5 10 20 30 40
E
SP EsP SxP
64 ** 28 28.2
14.1**** 34.4 38.8
21.9 37.6 33
F
47.7** 32.7 28.4
24.2* 32.2 33
28.1 35 38.6*
25 30 35 40 45

**Figure S7. Random forest model can predict the social stimulus in the SP task but not the specific stimulus among all six stimuli**

A. A color-coded confusion matrix of a multi-class Random Forest classifier using the changes in theta coherence during investigation bouts across the SP task to predict the type of investigated stimulus among the two possible stimuli (social and object).

B. As in A, for gamma coherence.

C. A color-coded confusion matrix of the same model using the changes in theta coherence during investigation bouts across all tasks to predict the type of investigated stimulus among the six possible stimuli.

D. As in C, for gamma coherence.

E. A color-coded confusion matrix for the Random Forest classifier employed for predicting the social context from $^{\Delta}\theta P$ values across all brain regions and stimuli. The scale of the accuracy's color code is shown to the right. The percentage of cases a label was predicted for each ground truth are marked in the middle of each spot. *$p<0.05$, **<0.01, ***<0.001, ****<0.0001, Mann-Whitney test, FDR corrected.

F. As in E, for the combination of $^{\Delta}\theta P$ and $^{\Delta}\theta Co$. *$p<0.05$, **<0.01, Mann-Whitney test, FDR corrected.

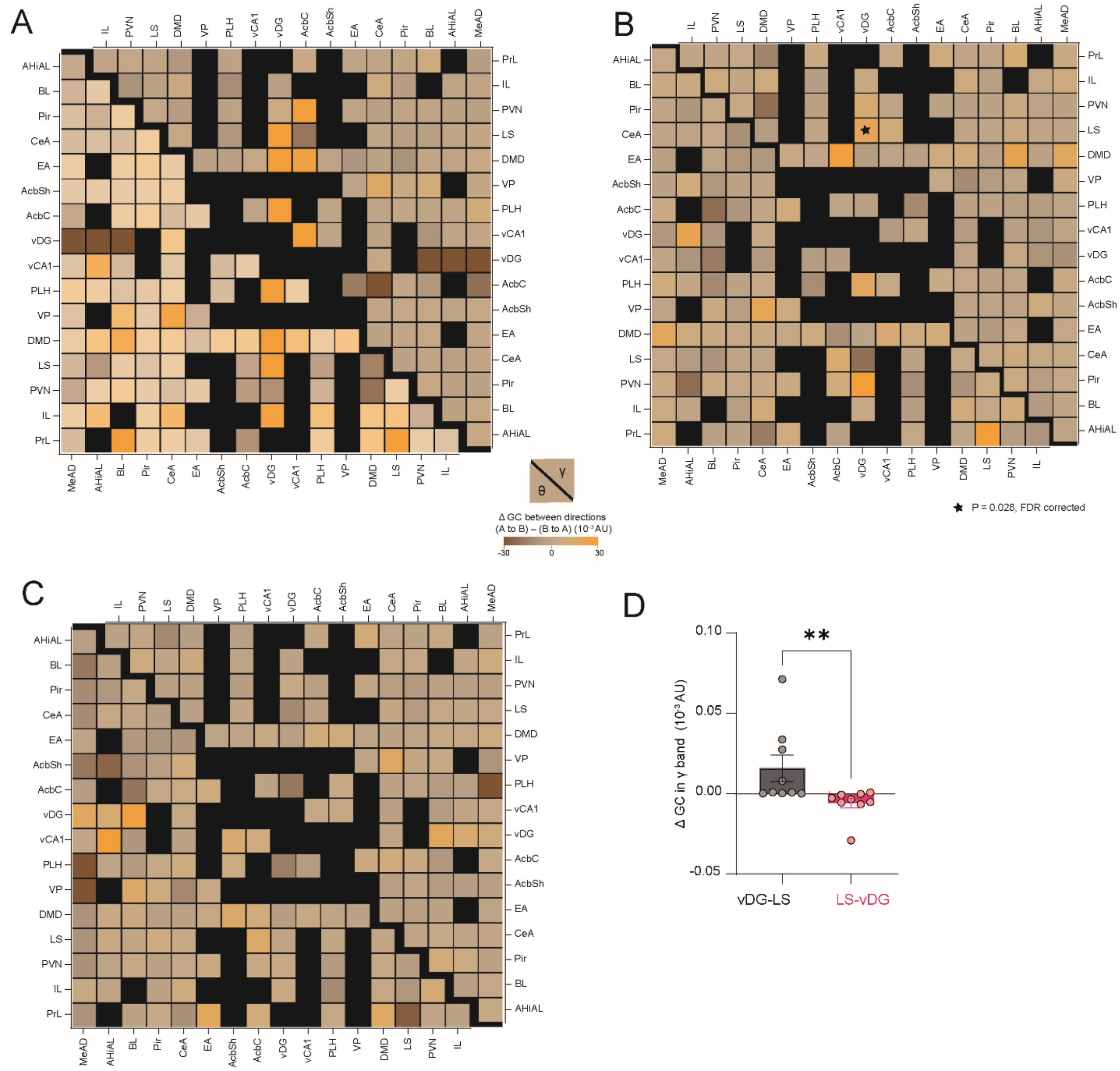


**Figure S8. Directionality of granger causality (GC) across coupled brain regions and tasks**

A. Color-coded matrices across all recorded brain regions of differences in GC changes (relative to baseline) during the SP task, between the two directions (region A to regions B and vice versa), for the theta (lower left) and gamma (upper right) bands.

B. As in A, for the EsP task. p<0.05, Mann-Whitney test, FDR corrected

C. As in A, for the SxP task.

D. Mean (±SEM) change in gamma GC from vDG to LS (Grey) and from LS to vDG pink. Wilcoxon matched pairs signed rank test, n = 9 sessions, W = -43, ***P*= 0.0078.

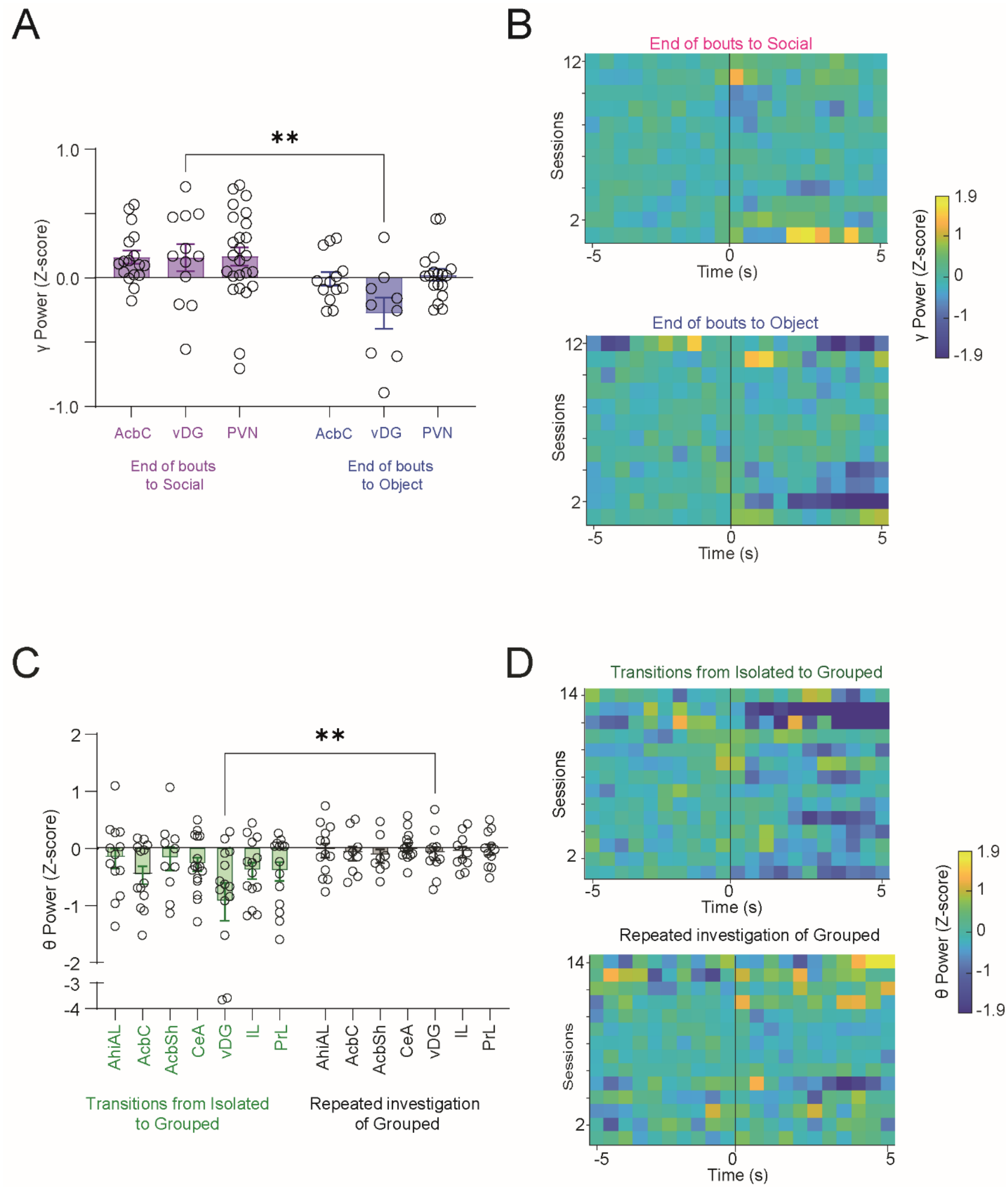


**Figure S9. Significant difference in gamma and theta power for specific behavioral events**

A. Mean (±SEM) Z-score values of gamma power change in the vDG at the end of long (>3 s) investigation bouts towards social (pink) and object (blue) stimuli during SP task sessions, shown for three brain regions where significant changes in GC were found in the gamma band.

B. Heat maps of male vDG gamma power changes before and during long (>3 s) investigation bouts towards the social (above) and object (below) stimuli across all SP task sessions. Each row represents the mean Z-score of all bouts in a single session (bins of 0.5s). Time point **'0'** represents the beginning of the bout. The color code is on the right.

C-D. As in A-B, for changes in theta power at the beginning of transitional (between stimuli) vs. repeated (for the same stimulus) investigation bouts of grouped social stimuli across EsP sessions.
